# A novel and highly effective mitochondrial uncoupling drug in T-cell leukemia

**DOI:** 10.1101/2020.06.23.168005

**Authors:** Victoria da Silva-Diz, Bin Cao, Olga Lancho, Eric Chiles, Amer Alasadi, Shirley Luo, David Augeri, Sonia Minuzzo, Stefano Indraccolo, Xiaoyang Su, Shengkan Jin, Daniel Herranz

## Abstract

T-cell acute lymphoblastic leukemia (T-ALL) is an aggressive hematologic malignancy. Despite recent advances in treatments with intensified chemotherapy regimens, relapse rates and associated morbidities remain high. In this context, metabolic dependencies have emerged as a druggable opportunity for the treatment of leukemia. Here, we tested the antileukemic effects of MB1-47, a newly developed mitochondrial uncoupling compound. MB1-47 treatment in T-ALL cells robustly inhibited cell proliferation via both cytostatic and cytotoxic effects as a result of compromised mitochondrial energy and macromolecule depletion, which severely impair nucleotide biosynthesis. Mechanistically, acute treatment with MB1-47 in primary leukemias promoted AMPK activation and downregulation of mTOR signaling, stalling anabolic pathways that support leukemic cell survival. Indeed, MB1-47 treatment in mice harboring murine NOTCH1-induced leukemias or human T-ALL PDXs led to a potent antileukemic effect with 2-fold extension in survival without overlapping toxicities. Overall, our findings demonstrate a critical role for mitochondrial oxidative phosphorylation in T-ALL and uncover MB1-47-driven mitochondrial uncoupling as a novel therapeutic strategy for the treatment of this disease.

## Introduction

T-ALL (T-cell acute lymphoblastic leukemia) is a relatively rare lymphoid neoplasm clinically characterized by elevated cell white counts, mediastinal thymic masses and frequent meningeal infiltration of the central nervous system (CNS)^1^. Management of newly diagnosed T-ALL cases is based on intensive chemotherapy regimens^2^. Despite the efficacy of these multiagent chemotherapy regimens, 20% of pediatric and over 50% of adult T-ALL cases still show primary resistance and/or relapse^3,4^, and the prognosis of these refractory/relapsed T-ALLs is extremely poor with a 5-year survival of less than 10%^5^. Remarkably, a high percentage of all casualties are due to toxic side effects during remission, usually from severe infections^6^. Additionally, most T-ALL survivors present further serious neurocognitive and cardiovascular impairments^7,8^, highlighting the need to develop safer strategies that might reduce morbidity and mortality rates.

A key feature of cancer cells is their capability to sustain chronic, uncontrolled proliferation and enhanced survival. In this context, one of the critical hallmarks of human cancer is cancer-specific metabolic rewiring, thus, selective targeting of primary cellular metabolic routes might be an attractive therapeutic option^9^. Indeed, antimetabolite drugs have been used in the clinic to treat hematologic malignancies for decades. One such example is aminopterin, to target “de novo” nucleotide biosynthesis^10^. Currently, the use of antimetabolites is widely extended in common regimes for leukemias, and include: purine antimetabolites, such as 6-mercaptopurine; pyrimidine antimetabolites, such as cytarabine; or anti-folates, such as methotrexate^11^. Notably, we recently demonstrated that targeting serine catabolism via SHMT inhibition shows highly antileukemic effects on its own and synergizes with methotrexate treatment both *in vitro* and *in vivo*, uncovering a promising novel pharmaceutical target for this disease^12^. Unfortunately, T-ALL cells can still develop mechanisms of resistance^13,14^ and antimetabolite drugs usually produce adverse effects in normal proliferative tissues like intestine, skin or bone marrow, causing severe diarrheas and/or immunosuppression^15,16^. In line with this, L-asparaginase, a crucial component for the success of conventional regimens in T-ALL cases, takes advantage of a particular metabolic vulnerability in leukemic cells^17,18^. The systemic administration of asparaginase dramatically depletes the plasma levels of asparagine by catalyzing de conversion of asparagine into aspartate and ammonia. Because asparagine is required for leukemia cell growth and T-ALL cells are highly dependent for extracellular pools of this amino acid, as they express low levels of asparagine synthetase (ASNS)^19^, this treatment shows an outstanding efficacy. However, adverse events occur in up to one third of patients, derived from hypersensitivity or liver dysfunction, resulting in chemotherapy cessation^20^. Thus, it is critical to design novel molecules with safer profiles to target metabolic routes on which leukemic cells exhibit higher dependency than normal proliferating cells do.

The detection of highly prevalent NOTCH1 activating mutations (∼60% of patients) in T-ALL ^21^ opened the opportunity for a more personalized targeted therapy and led to discovery of NOTCH1 signaling inhibitors, such as *γ*-secretase inhibitors (GSIs). GSIs block a critical proteolytical cleavage step required for NOTCH1 maturation and activation^22^ and are currently being explored in clinical trials for T-ALL relapsed/refractory cases. However, the responses observed as a single agent regimen have been limited due to the severe gastrointestinal toxicity^23^ and the acquisition of resistance mechanisms, such as the loss of PTEN expression^24^. These traits have limited the extensive use of these drugs, and current pre-clinical studies are focused on the discovery of new tumor liabilities to identify novel synthetic interactions. Interestingly, oncogenic NOTCH1 induces metabolic stress and promotes oxidative phosphorylation (OXPHOS) in T-ALL cells^25^ and the effects of NOTCH1 inhibition on central carbon metabolism are critical for its antileukemic activity in T-ALL^26^. Specifically, NOTCH1 inhibition translates into concomitant blocks in glycolysis and glutaminolysis, rendering T-ALL cells completely dependent on autophagy as a salvage pathway to survive^26^. In this context, we hypothesized that uncoupler drugs that target mitochondrial oxidative phosphorylation might be potent antileukemic agents. To test this hypothesis, we examined the efficacy and metabolic effects of a novel niclosamide-based second generation mitochondrial uncoupling drug in T-ALL cell lines, mouse primary leukemias and clinically relevant patient-derived xenografts *in vivo*.

## Methods

### Cell lines and culture conditions

Cells were cultured in standard conditions in a humidified atmosphere in 5% CO_2_ at 37°C in HyClone RPMI 1640 Media (SH3002701, Fisher Scientific) with 10-20% FBS (900-108, Gemini Bio-Products) and 100 U/mL penicillin and 100 μg/mL streptomycin (45000-652, VWR). DND41 (ACC 525), HPB-ALL (ACC 483), Jurkat (ACC 282), Molt3 (ACC 84), CCRF-CEM (ACC 240), TALL-1 (ACC 521) and Loucy (ACC 394) cells were obtained from Deutsche Sammlung von Mikroorganismen und Zellkulturen (DSMZ). The CUTLL1 cell line has been previously described^27^.

### Cell survival and cell size

To analyze cell survival, cells were plated in triplicates on 24 well plates: PTEN-positive cell lines were seeded at 300,000 cells per well in a final volume of 1 mL RPMI 1640 media under the experimental conditions described in each experiment; PTEN-negative cell lines were seeded at 100,000 cells per well; and non-NOTCH1 cell lines were plated at 400,000 cells per well, under the same culture conditions previously described. After 72 h of treatment, cells were counted using a Countess II FL instrument (Fisher Scientific) and values were normalized to their respective controls. Cell size changes induced by MB1-47 were estimated by flow cytometry (FSC-H) in G1-gated cells upon treatment for 72 h with MB1-47 or DMSO as a control.

### Oxygen Consumption Rate (OCR)

OCR rates were measured using a Seahorse Biosciences extracellular flux analyzer (XF24, Agilent Technologies) according to manufacturer’s instructions. Briefly, T-ALL cells were resuspended in Seahorse XF RPMI Medium (103576-100, Agilent Technologies) supplemented with 10 mM glucose (1003577-100, Agilent Technologies), 1 mM pyruvate (1003578-100, Agilent Technologies) and 2 mM glutamine (1003579-100, Agilent Technologies). 3-5 x 10^5^ cells per well were plated in XF24 Seahorse Biosciences plates pre-coated with Cell-Tak (354240, Corning) and spun down on the plate to ensure that cells were completely attached. OCR was analyzed by sequential injections of 1 μM oligomycin (O4876, Sigma), 1 μM FCCP (C2920, Sigma) or 2 μM MB1-47 (described in this study) and 0.5 μM rotenone (R8875, Sigma) in each well.

### In vivo efficacy of MB1-47 in NOTCH1-driven mouse T-ALLs

Animals were maintained in ventilated caging in specific pathogen-free facilities at New Brunswick RBHS Rutgers Campus. All animal housing, handling, and procedures involving mice were approved by Rutgers Institutional Animal Care and Use Committee (IACUC), in accordance with all relevant ethical regulations. Generation of Pten-conditional knockout NOTCH1-induced T-ALL tumors has been previously described ^26^. For survival studies, 1 x 10^6^ leukemia cells were transplanted from primary recipients into sub-lethally irradiated C57BL/6 (4.5 Gy) 6-8-week-old secondary recipients (Taconic Farms) by retro-orbital injection. To induce isogenic deletion of *Pten*, two days after leukemic cell transplantation, recipient mice were treated with 3 mg of tamoxifen (T5648, Sigma) dissolved in corn oil (AC405435000, ACROS Organics) at 30mg/mL via intraperitoneal injection. Subsequently, mice were divided randomly into two different groups: control group was fed with normal control diet (AIN-93M; Research Diets) while the treated groups were fed with AIN-93M diet containing MB1-47 (750 ppm, equivalent to a dose of 75 mg/kg b.w./day) and investigators were not blinded to group allocation. Animals were monitored for signs of distress or motor function at least twice daily, until they were terminally ill, whereupon they were euthanized.

To check for MB1-47 toxicity, MB1-47-treated healthy C57BL/6 mice were weighed every 2 days and sacrificed at day 30.

For MB1-47 acute treatment analyses in mouse primary leukemias, we injected leukemia cells into secondary recipients. We monitored mice until they presented clear leukemic signs with >60% GFP-positive leukemic cells in peripheral blood and mice were then treated twice (2 h apart) by oral gavage administration with control vehicle (0.5% carboxymethylcellulose) or MB1-47 (10 mg/Kg). 2 h after the last treatment, mice were euthanized, and leukemic spleens were collected for further analyses.

### Human primary xenografts

PDTALL#19 sample was provided by the University of Padova and has been previously described^26^. Patients were informed beforehand and signed the written consent. Samples were collected under the supervision of local Institutional Review Boards and analyzed under the supervision of Rutgers University Institutional Review Board.

### Measurement of hematological parameters

Blood from non-leukemic mice fed with control diet or MB1-47 containing diet (750 ppm) was collected in EDTA-containing tubes and was analyzed using an Element HT5 Veterinary Hematology Analyzer (Heska).

### Western blotting

Whole-cell extracts were prepared using standard procedures. After protein transfer, membranes were incubated with the antibodies anti-AMPK (1:1000, 5831S, Cell Signaling), anti-p-AMPK (1:1000, 2535S, Cell Signaling), anti-ACC (1:1000, 3676S, Cell Signaling), anti-p-ACC(1:1000, 11818S, Cell Signaling), anti-4E-BP1 (1:1000, 9644S, Cell Signaling), anti-p-4EB-P1 (1:1000, 2855S, Cell Signaling), and anti-β-actin-HRP (1:50.000; A3854, Sigma). Antibody binding was detected with a secondary antibody coupled to horseradish peroxidase (NA934; Sigma-Aldrich) using enhanced chemiluminescence (34578, Thermo Scientific).

### Flow cytometry analysis

To analyze thymic populations, single cell suspensions of total thymocytes were prepared by disrupting the tissues through a 40-μm filter. Red cells were removed by incubation with ammonium-chloride-potassium lysing buffer (155mM NH4Cl, 12mM KHCO3 and 0.1mM EDTA) for 5min at room temperature. Single cells were stained with anti-mouse fluorochrome-conjugated antibodies: CD4 PE-eFluor 610 (1:400, RM4-5, Thermo Fisher Scientific), CD8a PE (1:200, 53-6.7, BD Pharmingen), CD44 PerCPC-Cy5.5 (1:400, IM7, Thermo Fisher Scientific) and CD25 Alexa Fluor 488 (1:1000, 7D4, Thermo Fisher Scientific). (1:200, 53-6.7, BD Pharmingen). Total thymocytes were represented in a CD4 versus CD8a plot and CD4 SPs (CD4^+^CD8a^−^), CD8 SPs (CD4^−^CD8a^+^), CD4/CD8 DPs (CD4^+^CD8a^+^) and CD4/CD8 DNs (CD4^−^CD8a^−^) were gated. To differentiate discrete stages in early T-cell development, CD4/CD8 DNs were represented in a CD44 versus CD25 plot to characterize DN1 (CD44^+^CD25^−^), DN2 (CD44^+^CD25^+^), DN3 (CD44^−^CD25^+^) and DN4 (CD44^−^CD25^−^) populations, respectively. Cell cycle and apoptosis in T-ALL cells were performed after 72h of treatment with control (DMSO) or MB1-47 (2 – 4 μM). We used PI/RNase Staining Buffer (550825, BD Pharmingen) to analyze cell cycle distribution and we quantified apoptotic cells with PE-AnnexinV Apoptosis Detection Kit I (559763, BD Pharmingen).

To investigate the mitochondrial membrane potential, we performed TMRE staining (564696, BD Pharmingen). Briefly, T-ALL cells at a density of 1 × 10^6^ cells/mL were incubated in RPMI 1640 media with TMRE (100 nM) for 30 min at 37°C and 5% CO_2_. Dead cells were excluded using SYTOX™ Blue dead cell stain (S11348, Life Technologies).

All flow cytometry data were collected on an Attune NxT Flow Cytometer (ThermoFisher Scientific) and analyzed with FlowJo v10.6.2 software (BD).

### ^13^C labeling experiments

DND41 cells were pre-treated with DMSO or MB1-47 (4 μM) for 24 h in RPMI supplemented with 10 % dialyzed FBS. Then, cells were plated in triplicates in 6 well-dishes at 1.33 × 10^6^ cell/mL in RPMI supplemented with 10 % dialyzed FBS and, 10 mM U-^13^C-glucose or 4 mM U-^13^C-glutamine (Cambridge Isotope Laboratories), which included control (DMSO) or MB1-47 (4 μM). The isotopically labeled media was used for 4 h of incubation.

### Metabolite extraction

For T-ALL cells, cell suspensions were centrifuged (400 g, 2 min at RT). Media was removed by aspiration and the metabolome extraction was performed with 1 mL of methanol:acetonitrile:water (40:40:20) with 0.5% formic acid solution. After a 5 min of incubation on ice, acid was neutralized by the addition of 50 μL of 15 % ammonium bicarbonate. Finally, lysates were centrifuged (15,000 g, 10 min at 4 °C), and supernatants were frozen on dry ice and submitted for LC-MS analysis. For media, we diluted 20 μL of each respective media into 980 μL of ice cold solvent (40:40:20 methanol:acetonitrile:water +0.5% formic acid) followed by neutralization with 80 μL of 15 % ammonium bicarbonate. Samples were directly submitted to LC-MS analysis.

For leukemic spleens, leukemic mice were euthanized and spleens were quickly removed, minced and frozen. Subsequently, 20 to 30 mg per sample of frozen leukemic spleens were disrupted using a CryoMill at 20 Hz for 2 min. Pulverized samples were then mixed with methanol:acetonitrile:water (40:40:20) with 0.5% formic acid solution followed by 10 min incubation on ice and neutralization with 15 % ammonium bicarbonate. Finally, after centrifugation (14,000 g, 10 min at 4 °C), samples were transferred to clean tubes and sent for LC-MS analysis.

### LC-MS-based metabolomics

LC−MS analysis of the extracted metabolites was performed on a Q Exactive PLUS hybrid quadrupole-orbitrap mass spectrometer (ThermoFisher Scientific) coupled to hydrophilic interaction chromatography (HILIC). The LC separation was performed on Vanquish Horizon UHPLC system with an XBridge BEH Amide column (150 mm × 2.1 mm, 2.5 μM particle size, Waters, Milford, MA) with the corresponding XP VanGuard Cartridge. The liquid chromatography used a gradient of solvent A (95%:5% H_2_O:acetonitrile with 20mM ammonium acetate, 20mM ammonium hydroxide, pH 9.4), and solvent B (20%:80% H_2_O:acetonitrile with 20 mM ammonium acetate, 20 mM ammonium hydroxide, pH 9.4). The gradient was 0 min, 100% B; 3 min, 100% B; 3.2 min, 90% B; 6.2 min, 90% B; 6.5 min, 80% B; 10.5 min, 80% B; 10.7 min, 70% B; 13.5 min, 70% B; 13.7 min, 45% B; 16 min, 45% B; 16.5 min, 100% B. The flow rate was 300 μl/min. Injection volume was 5 μl and column temperature 25°C. The MS scans were in negative ion mode with a resolution of 70,000 at m/z 200. The automatic gain control (AGC) target was 3 × 10^6^ and the scan range was 75−1000. Metabolite features were extracted in MAVEN^28^ with the labeled isotope specified and a mass accuracy window of 5 ppm. For the ^13^C labeled samples, the isotope natural abundance and impurity of labeled substrate was corrected using AccuCor ^29^.

### Metabolic flux analyses

The metabolic flux analysis was performed using the elementary metabolite units (EMU) based method ^30^. The network contains 13 fluxes including the glutamate dehydrogenase flux fixed at 100. 5 fluxes were chosen as the free fluxes: pyruvate dehydrogenase flux, pyruvate carboxylase flux, ATP citrate lyase flux, isocitrate dehydrogenase exchange flux and fumarate hydratase exchange flux. The EMU model was constructed using R to calculate the metabolite labeling patterns when given the tracer labeling and the flux combination. The simulated labeling patterns for both U-^13^C-glucose and U-^13^C-glutamine tracers were compared to the measured labeling patterns for aconitate, α-ketoglutarate and malate. The global optimization by differential evolution was deployed to search for the best flux combination that fits the measured data the best.

### Statistical analysis

Statistical analyses were performed with Prism 8.0 (GaphPad). Unless otherwise indicated in figure legends, statistical significance between groups was calculated using an unpaired two-tailed Student’s t-test. Survival in mouse experiments was represented with Kaplan–Meier curves, and significance was estimated with the log-rank test.

## Results

### MB1-47 acts as a mitochondrial uncoupler in T-ALL cells

Mitochondria metabolism is critically involved in the control of bioenergetic and biosynthetic molecular pathways to sustain tumor cell survival and proliferation^31^. Specifically, previous studies showed that T-ALL cells rely on OXPHOS to maintain their proliferative capability^25^. Oxidative phosphorylation is coupled to protein complexes of the electronic transport chain (ETC) to transfer electrons from reducing equivalents NADH and FADH2 to oxygen, the final electron acceptor. As consequence of this electron flux, the ETC generates a high proton gradient across the mitochondrial inner membrane that is required to drive the ATP synthesis (Fig. 1a). Uncoupling drugs reduce this proton gradient and compromise the energy efficiency of mitochondria, leading to a futile oxidation of acetyl-CoA without generating ATP (Fig. 1a). Niclosamide (5-chloro-salicyl-(2-chloro-4-nitro) anilide) is an oral FDA-approved mitochondrial uncoupler for anthelmintic treatment. However, several studies indicate that niclosamide may have broad therapeutic applications for the management of more diverse diseases, including diabetes, viral infections or cancer^32^. Specifically, niclosamide presents a potent antiproliferative activity in a broad spectrum of cancer cell lines *in vitro*, being one of the top therapeutic candidates using the NCI 60 human cancer cell panel^33^ and its clinical potential is being assessed in ongoing clinical trials for prostate and colon cancer^34,35^. Notably, the action of niclosamide as mitochondrial uncoupler is well tolerated in normal cells^36,37^ and, used in its ethanolamine salt form (NEN), shows promising results in liver and metastatic colon cancer in mouse models *in vivo*^38,39^. However, both drugs have intrinsic limitations with transient and mild activities due to limited pharmacokinetic properties, hindering their use as clinical candidates. To improve the future translational applications for mitochondrial uncouplers, we synthesized a series of related niclosamide analogues, leading to the identification of MB1-47, in which one benzene ring is substituted by a benzothiazole group, while a methyl group is added to the other benzene ring. (Fig. 1b and Extended Data Fig. 1a). MB1-47 shows improved pharmacokinetic properties and increases the *in vivo* exposure of the drug ∼10-fold over NEN (Fig. 1c).

**Fig 1:**
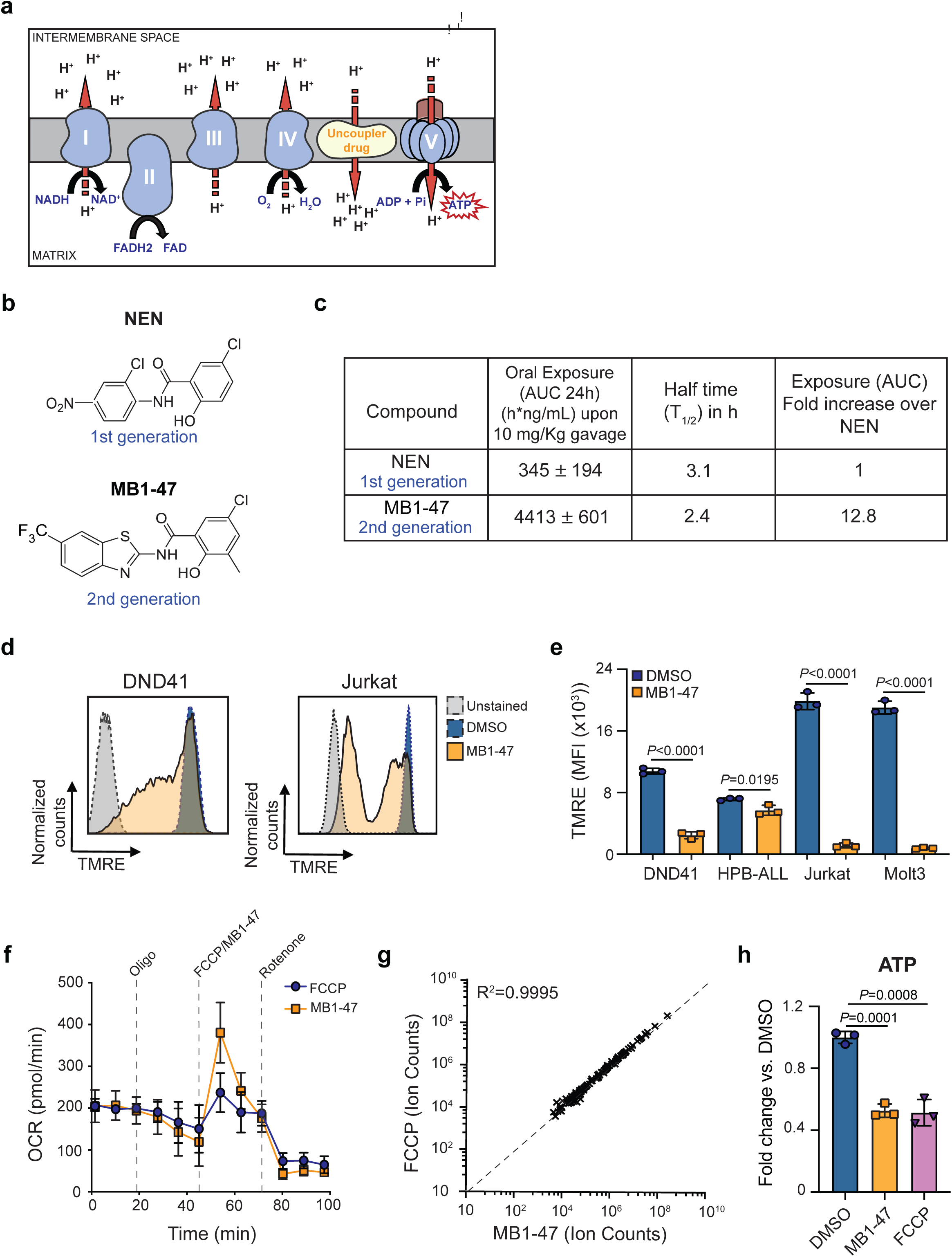
Chemical structure and mitochondrial uncoupling properties of MB1-47. **a**, Schematic illustration of the electron transport chain and the effects of uncoupling depolarizing the mitochondrial inner membrane. **b**, Chemical structure of NEN and MB1-47, niclosamide-based 1^st^ and 2^nd^ generation mitochondrial uncouplers, respectively. **c**, Pharmacokinetic/Pharmacodynamic properties of NEN and MB1-47 in mice *in vivo*. **d**, Representative flow cytometry histograms showing mitochondrial membrane potential measured by TMRE staining in live DND41 and Jurkat cells after 72 h of treatment with MB1-47 (4 μM and 2 μM, respectively). **e**, Geometric mean (± s.d.) quantification of TMRE staining from triplicates in two PTEN-positive (DND41 and HPB-ALL) and two PTEN-negative (Jurkat and Molt3) cell lines upon 72 h of MB-47 treatment. **f**, Oxygen consumption rate (OCR) in DND41 cells, under basal conditions or in response to the indicated mitochondrial inhibitors, measured in real time using a Seahorse XF24 instrument. Data are presented as mean ± s.d. of n = 4 wells. **g**, Metabolite levels in FCCP-treated *vs.* MB1-47-treated DND41 cells for 24 h (mean; n = 3). **h**, Relative quantification of ATP levels from DND41 triplicates treated with DMSO, FCCP or MB1-47 for 24 h (mean ± s.d.). Statistical significance (*P*) was determined by using unpaired two-tailed Student’s t-test. The goodness of fit (R^2^) was determined by using a simple linear regression model.

Next, we evaluated the uncoupling activity of MB1-47 in T-ALL cells, and we confirmed that MB1-47 treatment led to an expected and significant reduction in the mitochondrial membrane potential in all cell lines tested (Fig. 1d, e). In addition, similarly to FCCP (the gold-standard mitochondrial uncoupling drug), MB1-47 promoted an increase in oxygen consumption rate (OCR) even when ATP synthase is inhibited in presence of oligomycin (Fig. 1f). Moreover, metabolomic analyses of DND41 cells treated with MB1-47 or FCCP for 24 h revealed a very high correlation in the metabolic alterations in the presence of both drugs at their relative half maximal inhibitory constant (IC_50_) (Fig. 1g and Extended Data Fig.1b). These changes included a significant depletion in ATP levels (Fig. 1h), consistent with their uncoupling properties. Collectively, these data demonstrate that MB1-47 behaves as a mitochondrial uncoupling drug in T-ALL cells.

### MB1-47 treatment blocks T-ALL cell proliferation *in vitro*

To evaluate how leukemic cells respond to mitochondrial uncoupling, we treated with MB1-47 a panel of human T-ALL cell lines including both PTEN-positive^24^ (DND41, HPB-ALL and CUTLL1) and PTEN-negative^24^ (Jurkat, Molt3 and CCRF-CEM) cells, which are more resistant to glucocorticoids^40^ and to anti-NOTCH1 treatments^24^. These experiments revealed that MB1-47 has a potent intrinsic antileukemic activity in all cell lines tested, with an IC_50_ of ∼2-4 μM (Fig. 2a). Moreover, analysis of cellular sizes after MB1-47 treatment *in vitro* showed a marked reduction in cell diameters (Fig. 2b and Extended Data Fig. 2a), a common readout of antileukemic activity^24,26^. Finally, we observed similar antileukemic effects in non-NOTCH1 driven T-ALL cell lines (Loucy and TALL-1) (Extended Data Fig. 2b, c), overall suggesting that MB1-47 could be effective in a broad set of T-ALLs, independently of their oncogenic drivers and the mutational status of *NOTCH1* or *PTEN*. To dissect the antiproliferative effects of MB1-47, we next evaluated its impact on apoptosis and cell cycle progression in DND41 cells. These analyses revealed a combination of cytotoxic and cytostatic effects, as MB1-47 treatment results in markedly increased apoptosis (Fig. 2c, d) together with a simultaneous cell cycle arrest in G1 (Fig. 2e, f). These results were confirmed in a broader panel of T-ALL cell lines (Extended Data Fig. 2d, e). Altogether, our data demonstrate that MB1-47 has a strong intrinsic antileukemic activity *in vitro*.

**Fig. 2:**
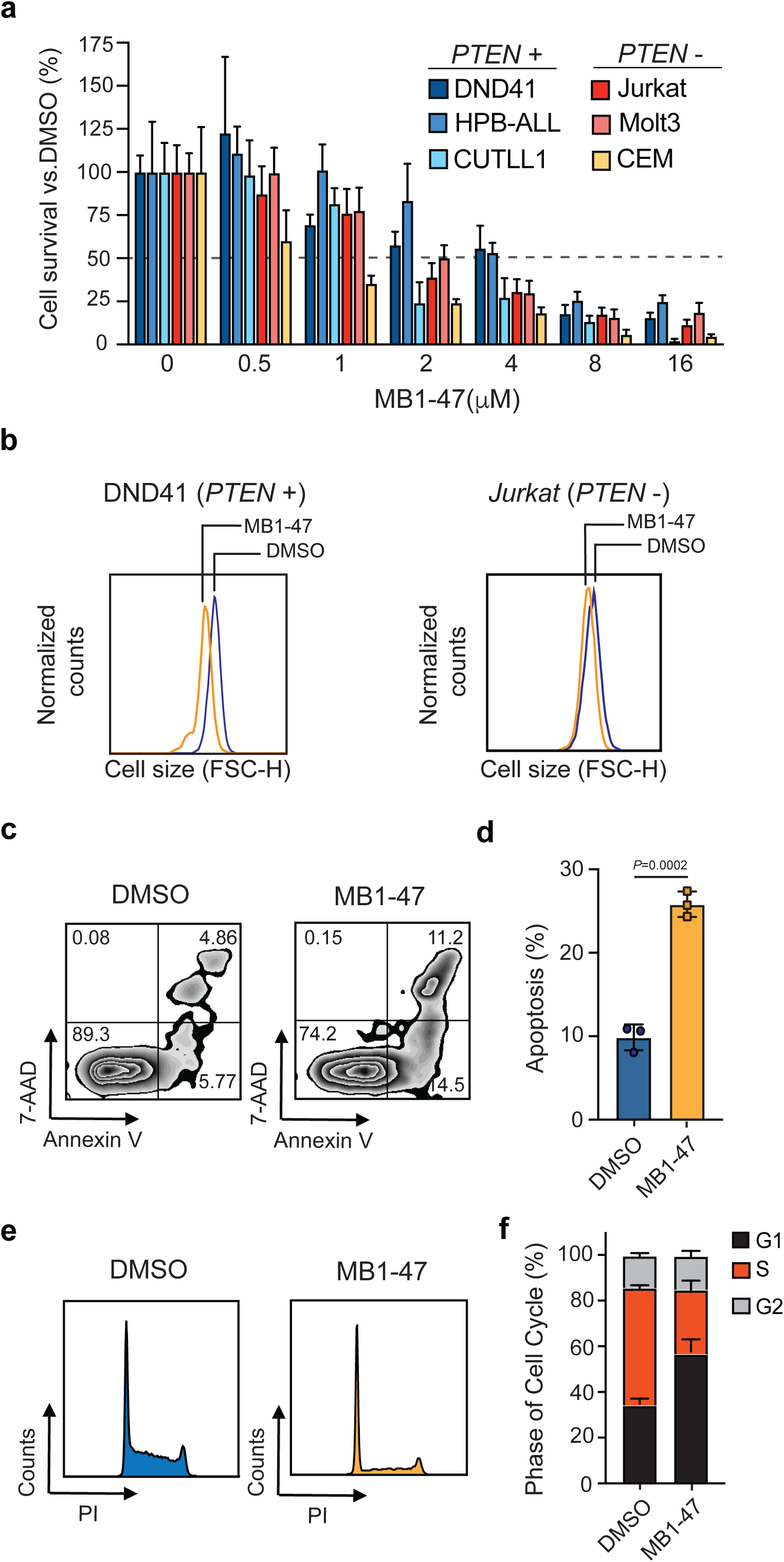
MB1-47 antileukemic effects in T-ALL cell lines *in vitro.* **a**, Relative cell survival of six independent human T-ALL cell lines (including PTEN-positive and PTEN-negative cells) in presence of MB1-47 at the indicated concentrations for 72 h (mean ± s.d.; n = 3). **b**, Representative flow cytometry histograms showing cell size changes in G1-gated DND41 and Jurkat cells treated with DMSO (control) or MB1-47 (4 μM) for 72 h. **c**, Representative flow cytometry plots of annexin V (apoptotic cells) and 7-AAD (dead cells) staining. Numbers in quadrants indicate percentage of cells. **d**, Quantification of apoptosis from DND41 triplicates treated for 72 h with DMSO (control) or 4 μM MB1-47 (mean ± s.d.). **e**, Flow cytometry representation of cell cycle analysis of DND41 cells treated with DMSO (control) or MB1-47 (4 μM) for 72 h. **f**, Quantification of cell cycle progression (mean ± s.d.) from triplicates from the indicated conditions as in **e**. Statistical significance (*P*) was determined by using two-tailed Student’s t-test.

### MB1-47 treatment alters the metabolic landscape of T-ALL cells *in vitro*

Although mitochondrial uncoupling drugs perturb oxidative phosphorylation-driven ATP production, their consequences on the global metabolic landscape of cancer cells are still unclear and their potential therapeutic benefit remains controversial. We first examined the bioenergetic status of MB1-47-treated leukemic cells. Previous reports indicated that, under normoxic conditions, myeloid leukemia cells cultured in presence of bone marrow derived mesenchymal stromal cells increase the expression of uncoupling protein 2 (UCP2), and these uncoupled cells accumulate lactate as result of reduced pyruvate oxidation without alteration in the glucose consumption^41^. Interestingly, exposure of T-ALL cells to MB1-47 for 24 h promoted increased glucose consumption (Fig. 3a) and lactate secretion (Fig. 3b) relative to untreated cells and this phenotype is common to all T-ALL cell lines evaluated (Extended Fig. 3a and Extended Fig. 3b), indicating a metabolic rewiring towards a highly glycolytic phenotype, as it has been previously described in non-small cell lung cancer (NSCLC) cells upon extended (> 24 h) metformin treatment^42^. To further dissect MB1-47 metabolic impact, we next performed untargeted liquid-chromatography mass spectrometry (LC-MS) analysis of intracellular water-soluble metabolites. Exposure to MB1-47 for 24 h caused profound alterations in the metabolic landscape of DND41 cells, leading to significant differences in 61 metabolites (Fig. 3c). Importantly, even if uncoupler drugs do not directly inhibit any component of the electronic transport chain, we noticed that MB1-47 decreased cellular NAD^+^/NADH and pyruvate/lactate ratios (Fig. 3d and Extended Fig. 3c), suggesting an electron acceptor insufficiency, similar to the metabolic perturbations caused by ETC inhibitors^43,44^. However, MB1-47 treated-T-ALL cells were unable to fully compensate this NAD^+^/NADH imbalance despite the increased lactate production (Fig. 3b). We next investigated the impact of MB1-47 on oxidative metabolism pathways. We observed TCA intermediates were among the most significantly altered metabolites (Extended Data Fig. 3d). Specifically, the levels of aconitate, citrate, α-ketoglutarate, glutamate, malate, fumarate and acetyl-CoA drop substantially upon MB1-47 exposure (Fig. 3e). Unexpectedly, we also observed a significant accumulation in the intracellular levels of succinate, similar to what is described in hypoxic ischemic tissues *in vivo*^45^. This could indicate either ETC exhaustion after prolonged exposure to uncoupler drugs or, alternatively, a defect in ETC complex II activity^46^. In addition, we detected significant changes in several amino acids (Fig. 3f and Extended Data Fig. 3d), with aspartate dropping ∼2.5-fold after MB1-47 treatment (Fig. 3g), and low aspartate levels could potentially limit leukemic cell proliferation under MB1-47 treatment, similar to what has been described for ETC inhibition in proliferating cells^43,44,47,48^. Since aspartate deficiency should impair *de novo* synthesis of nucleotides, we next quantified nucleotide pools and observed a significant reduction in the cellular levels of ATP, CTP, GTP and UTP and an accumulation of their metabolic precursors (IMP, AMP, CMP, GMP and UMP) (Fig. 3h and Extended Data Fig.3d). These results indicate that neither the aspartate deficiency nor the electron acceptor insufficiency caused the inhibition of nucleotide biosynthesis, as they are required for the conversion of IMP to AMP and GMP, respectively. However, the subsequent reactions for the final production of NTPs are catalyzed by ATP-dependent kinases. Thus, mitochondrial ATP could be the limiting metabolite for the normal cell cycle progression in T-ALL cells treated with MB1-47. Consistently with the increased UMP/UTP ratio (Fig. 3h), and given that UTP is required for the synthesis of UDP-sugars, we observed a significant depletion of UDP-glucose and UDP-N-acetyl-glucosamine (Fig. 3i), suggesting that the hexosamine biosynthetic pathway could also be required to support T-ALL cell growth. Most of these perturbations in nucleotide biosynthesis were common to a larger panel of T-ALL cell lines upon MB1-47 (Extended Fig. 3e-g). Taken together, our data show that nucleotide synthesis is severely compromised in the presence of MB1-47.

**Fig. 3:**
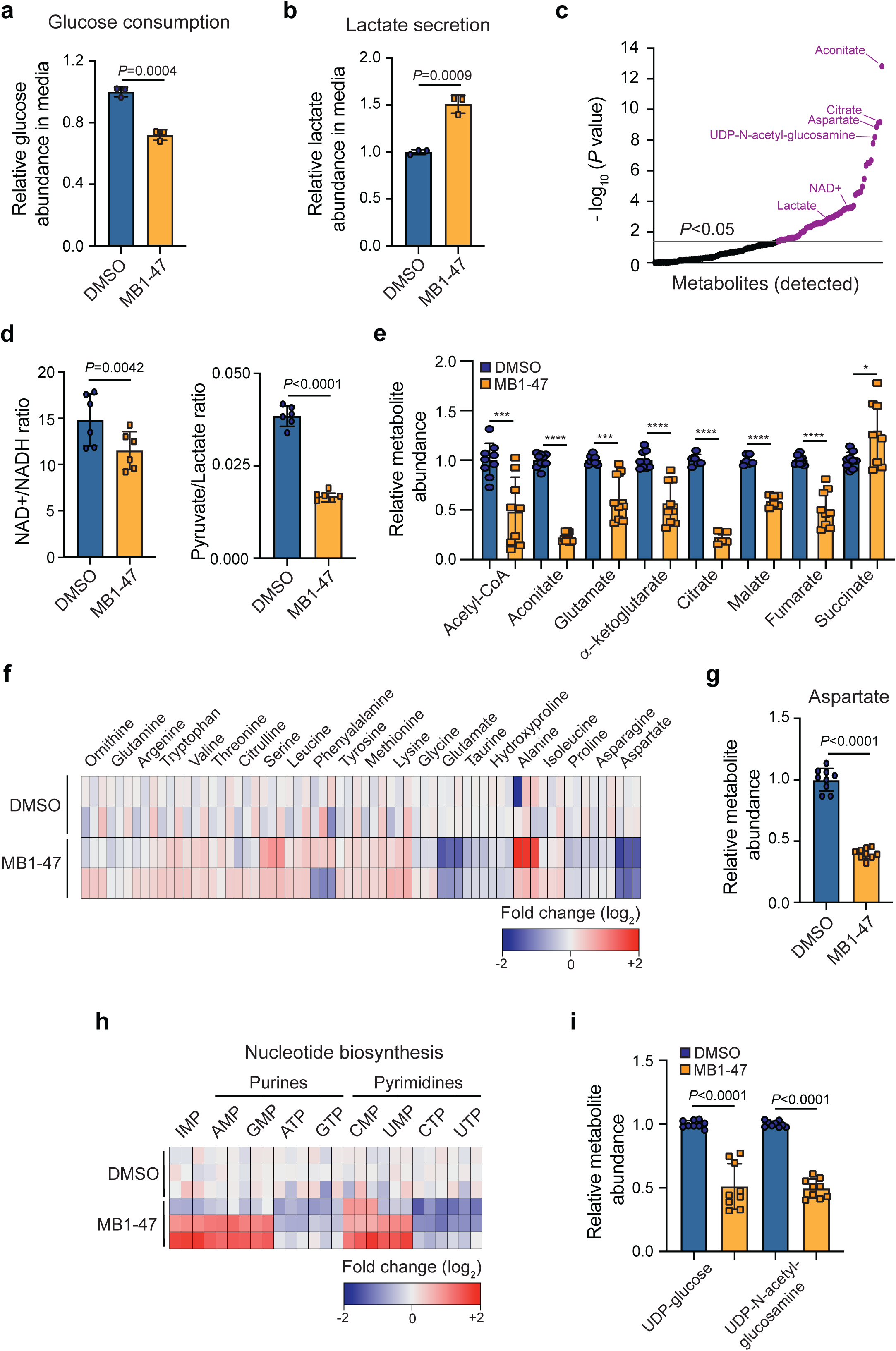
MB1-47 depletes TCA intermediates and NTPs in T-ALL cells. **a**, Relative glucose abundance in media from DND41 cells cultured in presence or absence of MB1-47 (mean ± s.d.; n=3). **b**, Relative lactate abundance in media from DND41 cells cultured in presence or absence of MB1-47 (mean ± s.d.; n=3). **c**, Significantly altered metabolites after MB1-47 exposure, ranked by *P* value (-log_10_ transformed). **d**, Intracellular ratio of NAD^+^/NADH (left) and pyruvate/lactate (right) in DND41 cells cultured in presence or absence of MB1-47 (n =6, from two independent replicates; bar graphs represent mean ± s.d.). **e**, Relative abundance of indicated TCA intermediates in DND41 cells cultured in presence or absence of MB1-47 (n = 9, from three independent experiments). **f**, Heat map showing differential intracellular amino acid abundances (log_2_) after MB-47 treatment, relative to DMSO-treated (control) cells. **g**, Relative abundance of aspartate in DND41 cells cultured in presence or absence of MB1-47 (n = 9, from three independent experiments; mean ± s.d.). **h**, Heat map showing differential intracellular nucleotide abundance (log_2_) after exposure to MB1-47, relative to DMSO-treated (control) cells. **i**, Relative abundance of indicated UDP-sugars in DND41 cells cultured in presence or absence of MB1-47. All measurements were determined after MB1-47 (4 μM) treatment for 24 h and are relative to DMSO-treated (control) cells. Statistical significance (*P*) was determined by using two-tailed Student’s t-test. * *P* < 0.05, ** *P* < 0.01, *** *P* < 0.001, **** *P* < 0.0001

### MB1-47 increases glucose flux and limits anapleurotic contributions to TCA cycle

Cancer cells upon acute pharmacological ETC inhibition use glutamine-dependent reductive carboxylation as the main pathway to maintain citrate levels^49^, and T-ALL cells are known to prominently rely on glutaminolysis to feed the TCA cycle^26^. To further understand MB1-47-driven effects on the TCA cycle, untreated or MB1-47 treated cells were cultured in medium containing uniformly labelled [U-^13^C]-glucose (Fig. 4a) or [U-^13^C]-glutamine (Fig. 4b), and ^13^C labeling of TCA intermediates was analyzed by LC-MS. Our data indicate that T-ALL cells increased the relative glycolytic flux to TCA in the presence of MB1-47 (Fig. 4c). Interestingly, examination of ^13^C isotopomer distribution after 4 h of labeling revealed that MB1-47 increased the fractional enrichments of m+4 aconitate and m+3 α-ketoglutarate (Fig. 4b), which can only be synthesized after a second round of the TCA cycle, suggesting an accelerated TCA cycle turnover. Consistently, [U-^13^C]-glutamine tracing experiments revealed an increment in the fractional enrichments of m+3 glutamate, m+3 α-ketoglutarate and m+2 malate (Fig. 4d). In addition, m+5 aconitate and m+3 malate (Fig. 4d) are reduced, suggesting that glutamine-dependent reductive carboxylation could be defective upon MB1-47 treatment. Moreover, MB1-47 reduced the flux through the glutamate dehydrogenase (73.53 in control cells vs. 60.92 in MB1-47-treated cells) (Fig. 4e), overall indicating a reduced contribution of glutamine into TCA cycle. Finally, these metabolic flux analyses upon MB1-47 treatment revealed increased citrate synthase (CS) activity (Fig. 4e), consistent the increased glycolytic flux (Fig. 4c), together with reduced pyruvate carboxylase (PC) and ATP-citrate lyase (ACLY) fluxes (Fig. 4e). Given that PC and ACLY are ATP-dependent limiting rate enzymes, our data suggest that mitochondrial ATP depletion upon MB1-47 treatment may limit anapleurotic contributions to TCA cycle.

**Fig. 4:**
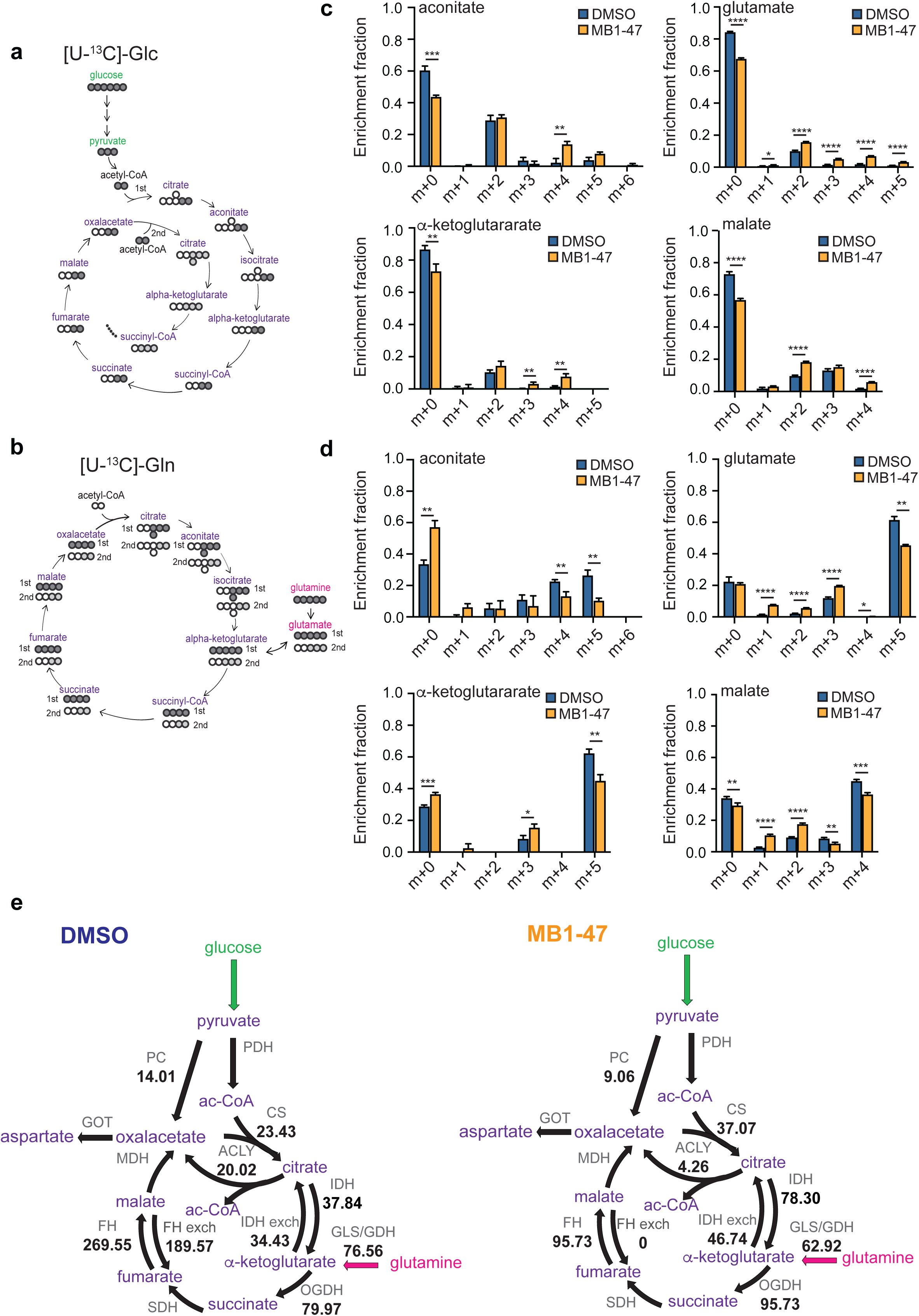
MB1-47 promotes increased glycolysis and glucose flux into TCA cycle. **a**, Schematic diagram of TCA cycle intermediates ^13^C labeling patterns in cells cultured with U-^13^C-glucose. **b**, Schematic diagram of TCA cycle intermediates ^13^C labeling patterns in cells cultured with U-^13^C-glutamine. **c**, Isotopologs for the indicated metabolites after 4 h of U-^13^C-glucose labeling in DND41 triplicates treated with MB1-47 or DMSO (control). **d**, Isotopologs for the indicated metabolites after 4 h of U-^13^C-glutamine labeling in DND41 triplicates treated with MB1-47 or DMSO (control). **e**, Relative flux activity of TCA cycle in untreated (DMSO) or MB1-47-treated DND41 cells. PC: Pyruvate carboxylase; PDH: Pyruvate dehydrogenase; CS: Citrate synthase; IDH: isocitrate dehydrogenase; GLS: Glutaminase; GDH: Glutamate dehydrogenase; OGDH: Oxoglutarate dehydrogenase; SDH: Succinate dehydrogenase; FH: Fumarate hydratase; MDH: Malate dehydrogenase; GOT: Glutamic oxaloacetic transaminase. Statistical significance (*P*) was determined by using multiple t-test. * *P* < 0.05, ** *P* < 0.01, *** *P* < 0.001, **** *P* < 0.0001

### MB1-47 treatment is well tolerated *in vivo*

To test the potential relevance of our findings *in vivo*, we first tested MB1-47 effects in healthy mice by feeding them with a control diet or a diet containing MB1-47 (750 ppm). First, we analyzed potential drug-induced toxicities upon long-term exposure to MB1-47 for over 30 consecutive days. Importantly, MB1-47 treatment is well tolerated, as revealed by the unchanged mouse weight during the entire experiment (Fig. 5a). Moreover, mice did not present any sign of anemia or other hematologic alterations (Fig. 5b and Extended Data Fig. 4a). Furthermore, detailed immuno-phenotypic analyses of T-cell development in these mice revealed no significant differences in the percentages of different thymocyte populations (Fig. 5c-e), including CD4/CD8-double negative (DN1, DN2, DN3 and DN4) (Fig. 5e, f), CD4/CD8-double positive (Fig. 5c, d) or single positive CD4 or CD8 cells (Fig. 5c, d). Therefore, our data suggest that there might be a good therapeutic window for the use of MB1-47 *in vivo*.

**Fig. 5:**
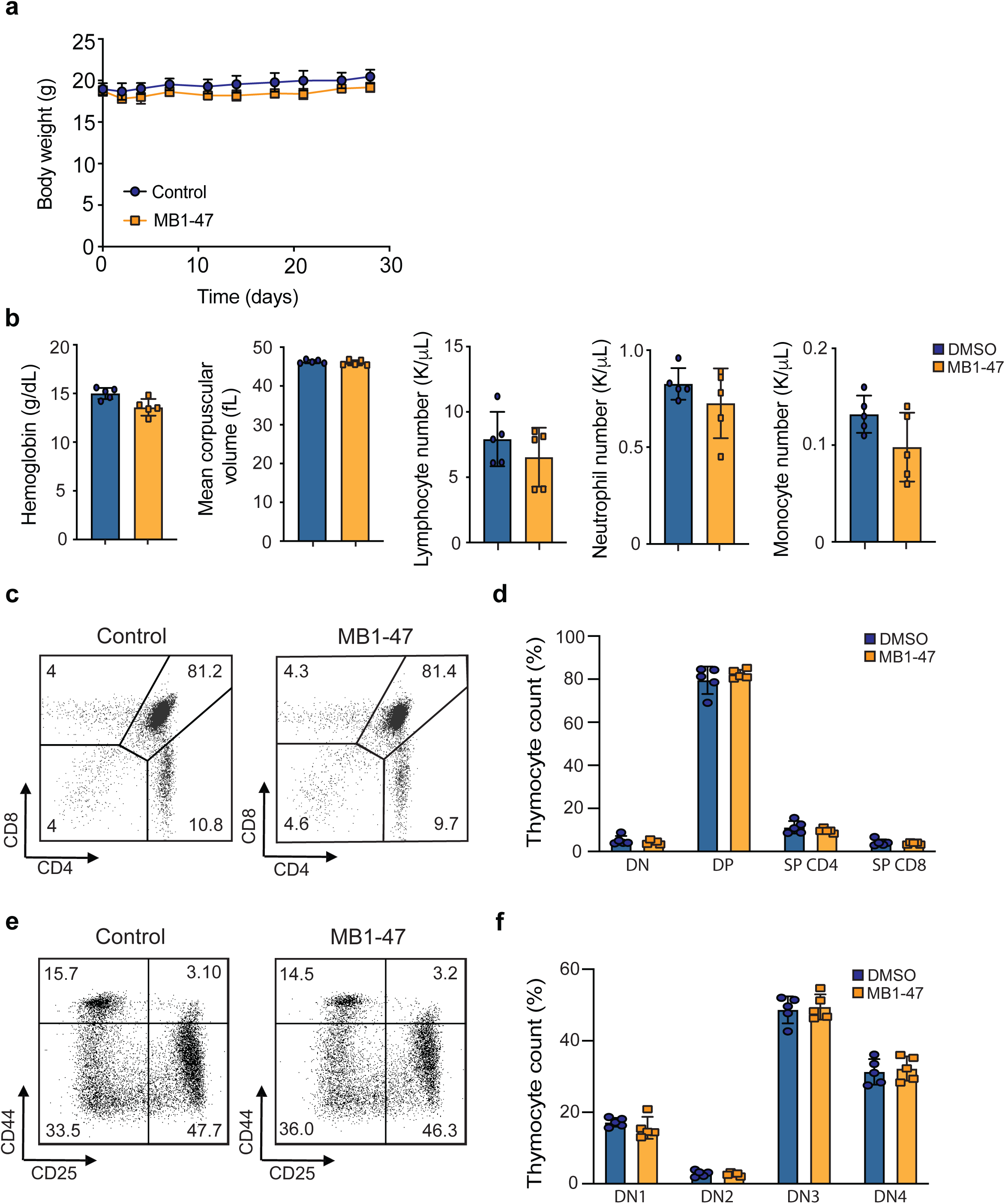
MB1-47 effects in healthy mice *in vivo*. **a**, Body weight of mice fed with control diet or MB1-47 diet evaluated during 30 consecutive days (mean ± s.d.; n = 5). **b**, Hematological parameters in mice fed with control diet or MB1-47 diet (mean ± s.d.; n = 5). **c**, Representative flow cytometry plots of thymic populations upon anti-CD4 and CD8a staining of thymocytes from mice fed with control diet or MB1-47 diet. **d**, Immunophenotypic quantification of the indicated thymocyte populations (mean ± s.d.; n = 5). **e**, Representative flow cytometry plots of the different stages of early T-cell development (DN1-DN4) in mice fed with control diet or MB1-47 diet. **f**, Quantification of different DN thymocyte stages (mean ± s.d., n = 5). All parameters were evaluated after 30 consecutive days of treatment. Statistical significance (*P*) was determined by using two-tailed Student’s t-test.

### Acute MB1-47 treatment in T-ALL *in vivo* reveals an altered metabolic profile

Next, we investigated the metabolic impact of MB1-47 in T-ALL *in vivo* taking advantage of our previously established NOTCH1-induced mouse primary leukemias^26^. Mice showing signs of overt leukemia (> 60% GFP+ leukemic cells in peripheral blood) were exposed to MB1-47 or vehicle as a control over a 4 h-period (Fig. 6a and Methods), followed by untargeted (LC-MS) metabolomic analyses of the untreated or MB-47-treated leukemic spleens. Even after this short time of exposure, we detected significant differences in 20 metabolites between untreated and MB1-47-treated leukemias (Fig. 6b). Remarkably, ∼60% of the detectable metabolites were also significantly altered after the MB1-47 treatment *in vitro*, indicating that MB1-47 promotes a similar metabolic impact *in vivo*. Indeed, we also observed a slight reduction in the pyruvate/lactate ratio (Fig. 6c), despite the potential interference from non-leukemic stromal cells. Remarkably, and consistent with the metabolic profiles of MB1-47-treated T-ALL cell lines *in vitro* described previously, we detected a significant reduction in the intratumor levels of aspartate, glutamate, and malate (Fig. 6d), indicating that these metabolic perturbations constitute the core of the immediate response to uncoupler drugs and may mediate the first steps in the metabolic cascade for its antiproliferative activity. To dissect the molecular mechanism behind this metabolic stress, we evaluated the status of AMP-activated serine/threonine protein kinase (AMPK) in primary leukemias upon treatment with MB1-47. AMPK is activated under conditions of energy stress such as nutrient deprivation or hypoxia and, through the inhibition of mTOR, acts a metabolic checkpoint suppressing anabolic biosynthetic processes^50^. Our results show that MB1-47 promotes increased phosphorylation of AMPKα (Fig. 6e) with concomitant phosphorylation of its downstream effector ACC (acetyl-CoA carboxylase) (Fig. 6e). In line with this, we also observed lower levels of phosphorylated 4E-BP1, supporting decreased mTOR activity (Fig. 6e). Previous reports demonstrated that NOTCH1 regulates mTOR activity in T-ALL cells^51^ and mTORC2 depletion increases mouse survival in NOTCH1-driven leukemic models^52^. Thus, our data suggest that the MB1-47-promoted metabolic stress is sensed by AMPK, driving downregulation mTOR, hampering anabolic pathways and restricting cell cycle progression.

**Fig 6:**
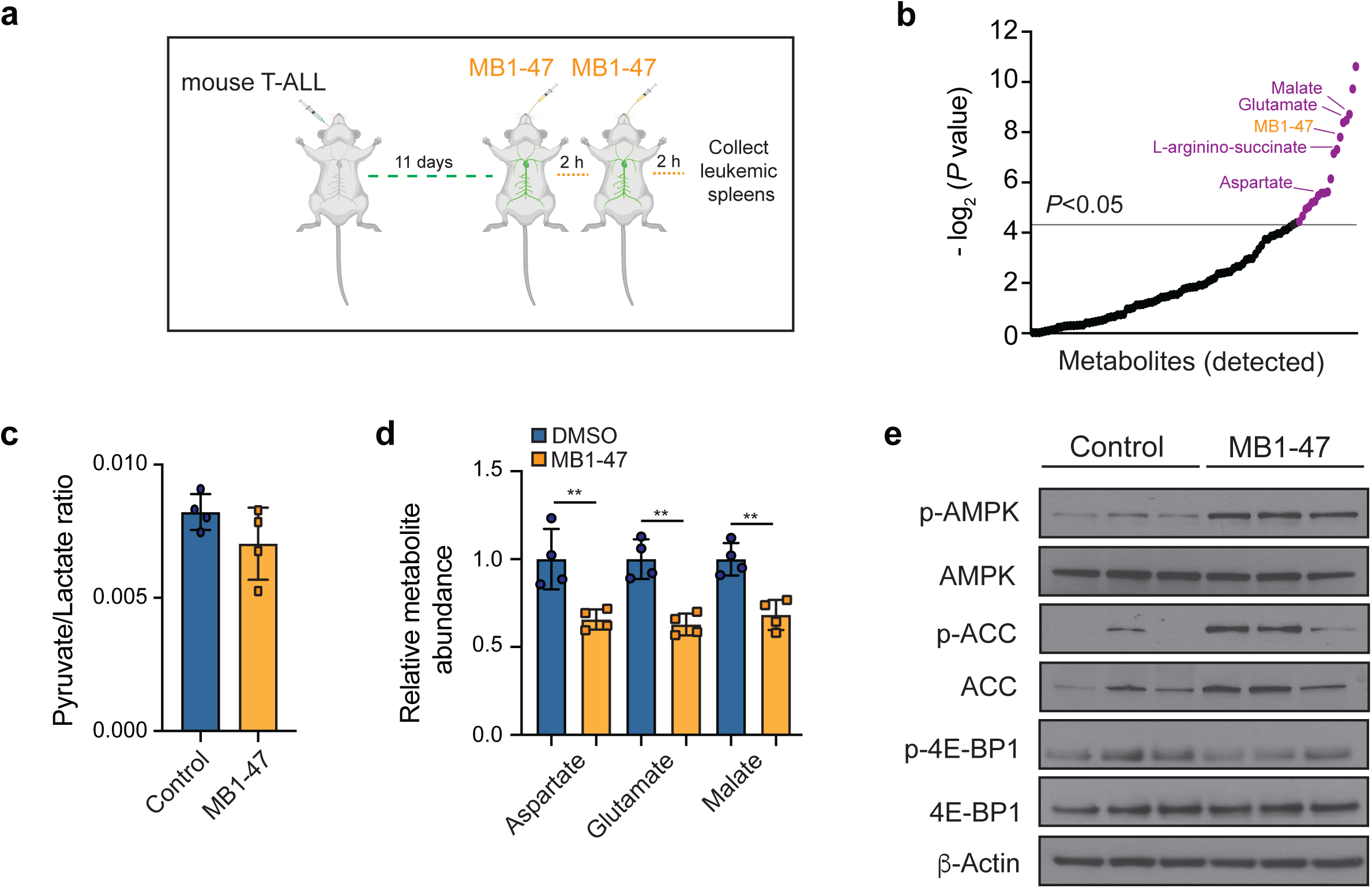
Metabolic and signaling effects upon acute MB1-47 treatment of leukemic mice *in vivo*. **a**, Schematic illustration of acute exposure to MB1-47 *in vivo* (detailed in Methods). **b**, Significantly altered metabolites after acute exposure to MB1-47, ranked by *P* value (-log_2_ transformed). **c**, Ratio of pyruvate/lactate abundances from leukemic spleens after acute treatment with Vehicle (control) or MB1-47 (mean ± s.d.). **d**, Relative abundance of the indicated metabolites in leukemic spleens after acute treatment with MB1-47 *in vivo*. **e**, Immunoblot analyses of AMPK, ACC and 4E-BP1 in leukemic spleens from mice acutely treated with Vehicle (control) or MB1-47 for 4 h. Statistical significance (*P*) was determined by using two-tailed Student’s t-test. ** *P* < 0.01, *** *P* < 0.001.

### MB1-47 shows antileukemic effects in mouse primary T-ALL and patient derived xenografts *in vivo*

Since MB1-47 showed a favorable safety profile together with promising antileukemic effects *in vitro*, we next examined whether MB1-47 could impair tumor growth in mice harboring NOTCH1-induced primary leukemias *in vivo*. Here, we took advantage of our previously established *Pten*-conditional knockout (Rosa26^Cre-ERT2/+^Pten^f/f^) isogenic NOTCH1-induced leukemias^26^ in order to analyze the effects of MB1-47 on the survival of mice harboring either Pten-positive or Pten-deficient (upon tamoxifen-induced activation of the Cre recombinase) leukemias. In this context, leukemic mice were randomly assigned into four groups, in which mice were fed with control diet or diet containing MB1-47, and treated with corn oil or tamoxifen, in order to induce *Pten* deletion. Remarkably, and consistent with our previous *in vitro* results, MB1-47 treatment translated into very drastic antileukemic effects *in vivo* with significantly extended survival (∼2-fold) of mice harboring Pten-positive leukemias (Fig. 7a, d; median survival in control mice = 14 days, median survival in MB1-47 treated mice = 28 days). In addition, and most notably, this antileukemic effects were also maintained in mice harboring Pten-deficient leukemias (Fig. 7b, d). Despite the expected acceleration of disease kinetics upon Pten loss^26^ (median survival in control mice = 11 days, Fig.7b d), MB1-47 treatment still conferred a similar ∼2-fold increase in disease latency in mice harboring Pten-null leukemias (median survival in MB1-47 treated mice = 21 days, Fig. 7b, d). These findings are clinically-relevant, as loss of Pten is known to mediate resistance to glucocorticoid treatment^40^, as well as resistance to anti-NOTCH1 therapies^24,26^. Finally, we also tested the therapeutic efficacy of MB1-47 in mice harboring a patient derived T-ALL xenograft (PDX) harboring a mutation in NOTCH1 (PDTALL#19). In this context, treatment with MB1-47 translated into a similar ∼2-fold increase in survival (Fig. 7c, d). Altogether, our data demonstrate that disrupting oxidative phosphorylation using uncoupler drugs limits leukemia progression and, more specifically, MB1-47 treatment could be a novel promising therapeutic strategy for T-ALL patients.

**Fig 7:**
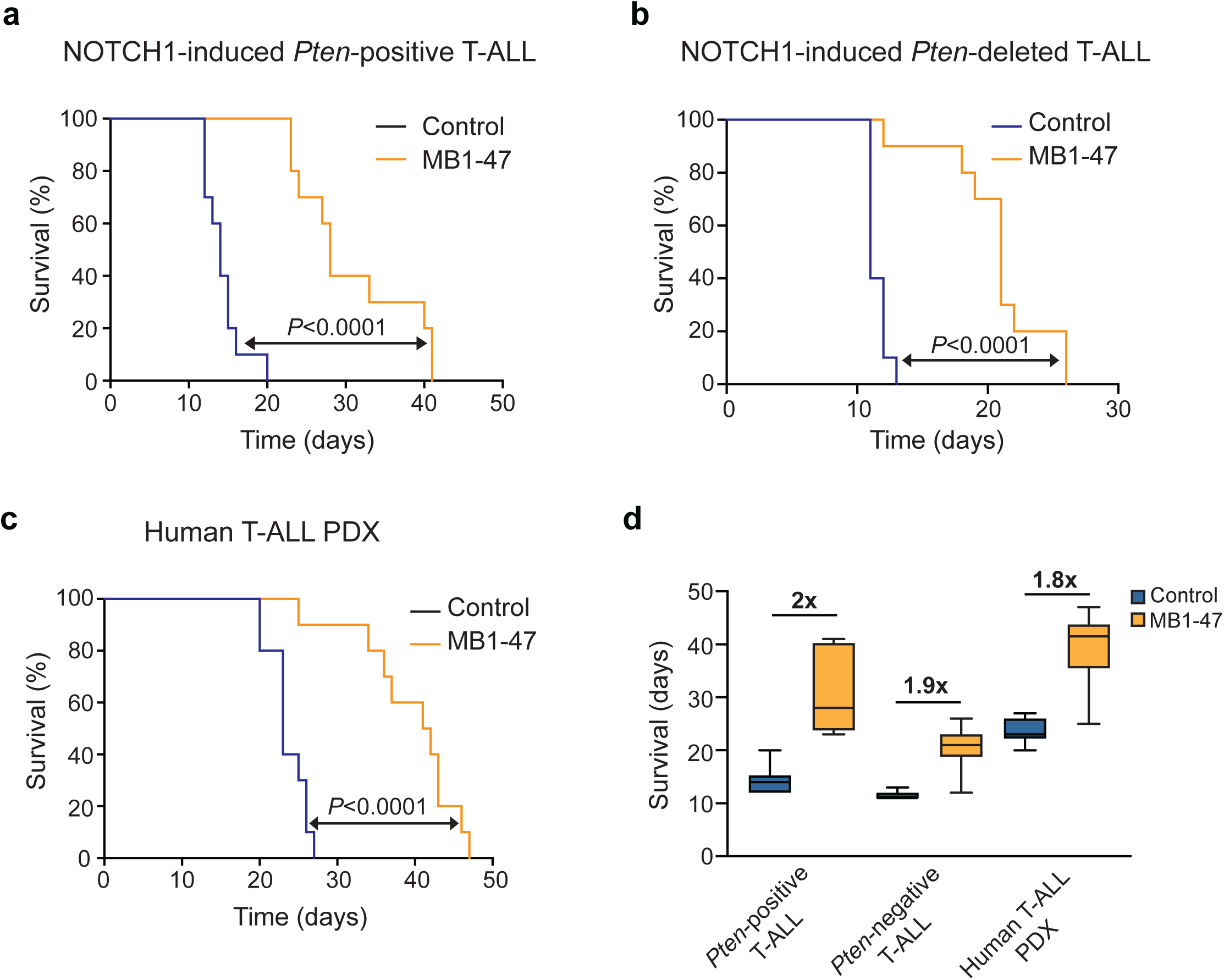
MB1-47 shows antileukemic effects in mouse primary leukemia and human T-ALL PDXs *in vivo*. **a-b**, Kaplan-Meier survival curves of mice harboring isogenic *Pten*-positive (**a**) and *Pten*-deleted (**b**) T-ALLs treated with MB1-47-containing diet or control diet. **c**, Kaplan-Meier survival curves of mice harboring a human T-ALL PDX treated with control diet or MB1-47-containing diet. **d**, Boxes represent the median and the first and third quartiles of survival, and whiskers indicate the minimum and maximum of all data values. *P* values were calculated with the log-rank test; n = 10 mice per group.

## Discussion

The development of new therapeutic agents for the treatment of T-ALL remains an unmet clinical need. Recent advances have improved our understanding of the basic biology and genetics driving this disease^53^, however, clinical management of T-ALL cases is still based on intensive salvage chemotherapy regimens. Even if these treatments have considerably improved clinical outcomes in the last decades^3^, many patients still relapse and/or present long-term adverse health impairments, highlighting the urgent need to discover more effective interventions with less overlapping toxicities. Oncogenic transformation, driven by MYC, RAS, NOTCH or loss of PTEN, is closely linked to leukemia-associated metabolic rewiring and these oncogene-driven metabolic dependencies may offer a more druggable opportunity for translational purposes^11^. For decades, the cancer metabolism field focused efforts on perturbing the enhanced glycolysis (Warburg effect) observed in cancer cells, however, recent studies demonstrated that mitochondrial metabolism is essential for tumorigenesis^54-57^. In the present study, we developed and characterized MB1-47, a novel niclosamide-based second generation mitochondrial uncoupling drug and we provide evidence that T-ALL cells are highly dependent on OXPHOS and mitochondrial ATP for proliferation and survival. Our *in vitro* data suggest that MB1-47 could be effective in different T-ALL subtypes, independently of oncogenic drivers and mutational status of *NOTCH1* and *PTEN*. Analyses of global metabolic profiling revealed that MB1-47 treatment generates an environment of energy deficiency and macromolecule depletion that severely compromises nucleotide biosynthesis, leading to apoptosis and cell cycle arrest, comparable to what has been observed after the inhibition of complex I in AML^58^. Even if MB1-47 does not directly inhibit any ETC complex, we observed a low NAD^+^/NADH ratio, similar to the metabolic phenotype in proliferating cells with impaired mitochondrial respiration^43,44^. Given that MB1-47 promotes increased glucose consumption and lactate secretion, it is reasonable to speculate that this glycolytic phenotype may be a compensation mechanism for an electron acceptor insufficiency^43^. Beyond this NAD^+^ deficiency, we found that MB1-47 also reduces the levels of most TCA intermediates except succinate. The intracellular accumulation of succinate may indicate defects in the complex II of the ETC, as it has been previously described after venetoclax plus azacytidine in AML leukemic stem cells^46^. Alternatively, the increase in succinate could be due to hypoxia conditions caused by ETC exhaustion after prolonged treatment with MB1-47, mimicking the succinate accumulation observed in ischemic tissues^45^. However, the importance of succinate accumulation or whether reduction of its levels could ameliorate the metabolic phenotype after MB1-47 treatment remain to be elucidated. In addition, MB1-47 led to significant changes in several amino acids and, in concordance with the observed electron acceptor insufficiency, aspartate synthesis seems to be limited^43,44^. A lingering question here is how a mitochondrial uncoupler creates a metabolic phenotype with high correlation to the metabolic perturbations previously described for inhibition of complex I and II of the ETC^43,44,46-48,58^. However, our data indicate that the aspartate deficiency or electron acceptor insufficiency cannot fully explain the compromised nucleotide biosynthesis after MB1-47 exposure and suggest that mitochondrial ATP might be the limiting metabolite. Related to this, a recent study demonstrated mitochondrial ATP, but not glycolytic ATP, regulates fatty acid uptake and transport^59^.

Moreover, metabolic isotope-labeling analyses indicate that mitochondrial ATP depletion may limit the anapleurotic contributions to TCA cycle, slowing down the metabolic fluxes through PC and ACLY, two ATP-dependent rate-limiting enzymes. Indeed, in normal conditions, cells use pyruvate carboxylation to produce aspartate, and the decreased flux through PC may explain in part the reduced levels for this amino acid. Furthermore, PC deficiency also results in depletion of TCA cycle intermediates and acetyl-CoA, two phenotypes induced by MB1-47. Acetyl-CoA is a crucial metabolite for energy production, lipid metabolism and epigenetic modifications and it is used for the synthesis of important molecules, such as cholesterol and UDP-sugars^60,61^. The substantial drop in acetyl-CoA after MB1-47 treatment could be explained in part to the limited ACLY flux and may contribute to the dramatic UDP-N-acetylglucosamine depletion. Furthermore, previous studies revealed a prominent role for glutaminolysis to fuel TCA-cycle downstream of NOTCH1^26^. Genetic inactivation or pharmacological inhibition of glutaminase strongly synergized with anti-NOTCH therapies and exhibited marked anti-T-ALL activity *in vivo*^26^. In line with this, MB1-47 treatment in T-ALL PTEN-positive cell lines promoted an accelerated TCA turnover and decreased glutamine-derived carbon input to the TCA cycle, indicating that the limited glutamine input may contribute to the enhanced antileukemic activity of this drug. We also demonstrate that MB1-47 is selectively toxic against leukemic cells without affecting T-cell development in the thymus or other hematologic populations in peripheral blood from healthy mice, thus, MB1-47 shows an acceptable safety profile.

In addition, acute exposure of leukemic mice to MB1-47 led to depletion of some TCA intermediates, such as malate and aspartate, suggesting that the observed metabolic shutdown *in vitro* might be reliably translated *in vivo*. A recent report indicates that AMPKα1 can act as cell-autonomous tumor suppressor and mediate the therapeutic effect of phenformin in T-ALL development^62^. In contrast, Kishton et al. showed that AMPK deficiency reduced disease burden and pointed that AMPK switches to function as tumor promoter during T-ALL progression^25^. Our data indicate that MB1-47 treatment *in vivo* promotes metabolic stress, activates AMPKα and downmodulates mTOR pathway, restricting anabolic pathways that support leukemic cell survival. However, the role of AMPKα−mTOR pathway mediating the T-ALL responses to MB1-47 remains to be fully elucidated.

Importantly, MB1-47 exhibited strong antileukemic activity as a single agent and significantly extended the survival of mice harboring Pten-positive leukemias. Strikingly, and unlike glutaminase inhibition^26^, MB1-47 as single agent also delayed T-ALL progression in mice harboring Pten-null leukemias. Our results uncover a mitochondrial respiration dependency for Pten-null leukemias, despite the fact that Pten loss induces a hyperglycolytic phenotype that drives resistance to NOTCH inhibitors^24,26^. In addition, our findings suggest that mitochondrial uncoupling could be an attractive therapeutic strategy in cases with PTEN loss, which are known to be resistant to glucocorticoid treatment^40^ or anti-NOTCH1 therapies^26^. Finally, MB1-47 showed a very similar therapeutic effect in a human PDX-T-ALL model. Overall, our findings suggest a critical role for mitochondrial oxidative phosphorylation in T-ALL and demonstrate that the novel MB1-47 mitochondrial uncoupling drug could be an attractive therapeutic strategy for the treatment of T-ALL patients.

## Acknowledgements

We thank all members of the Herranz Laboratory for helpful discussions. Work in the laboratory of D. H. is supported by the US National Institutes of Health (R01CA236936), a Research Scholar Grant from the American Cancer Society (RSG-19-161-01-TBE), the Alex’s Lemonade Stand Foundation, the Leukemia Research Foundation, the Children’s Leukemia Research Association and the Gabrielle’s Angel Foundation for Cancer Research. V.da Silva-Diz is funded by the New Jersey Commission on Cancer Research (DCHS19PPC008). In addition, Rutgers Cancer Institute of New Jersey shared resources supported in part by the National Cancer Institute Cancer Center Support Grant P30CA072720 were instrumental for this project and, more specifically, the Metabolomics Shared Resource (P30CA072720-5923).

## Author contributions

V. dS. performed most molecular biology, cellular and animal experiments and wrote the manuscript. O.L. assisted with cellular experiments. S.L. assisted some animal experiments. E.C. processed metabolomic samples. B.C., D.A. designed and synthesized MB1-47. A.A. did initial mitochondrial uncoupling characterization of MB1-47. X.S. supervised and analyzed all the metabolomic experiments. S.M. and S.I. generated and provided the human primary xenograft used in this paper. S.J. conceived the idea, assisted with design and synthesis of MB1-47 and assisted with design of study. D.H. conceived the idea, designed the study, supervised the research and wrote the manuscript with V. dS.

## Author information

B.C., D.A., and S.J. are co-inventors of the patent covering MB1-47. S.J. is a co-founder of Mito Biopharma which licensed the patent.

The rest of the authors declare no competing financial interests.

## Extended Figures

**Extended Fig. 1:**
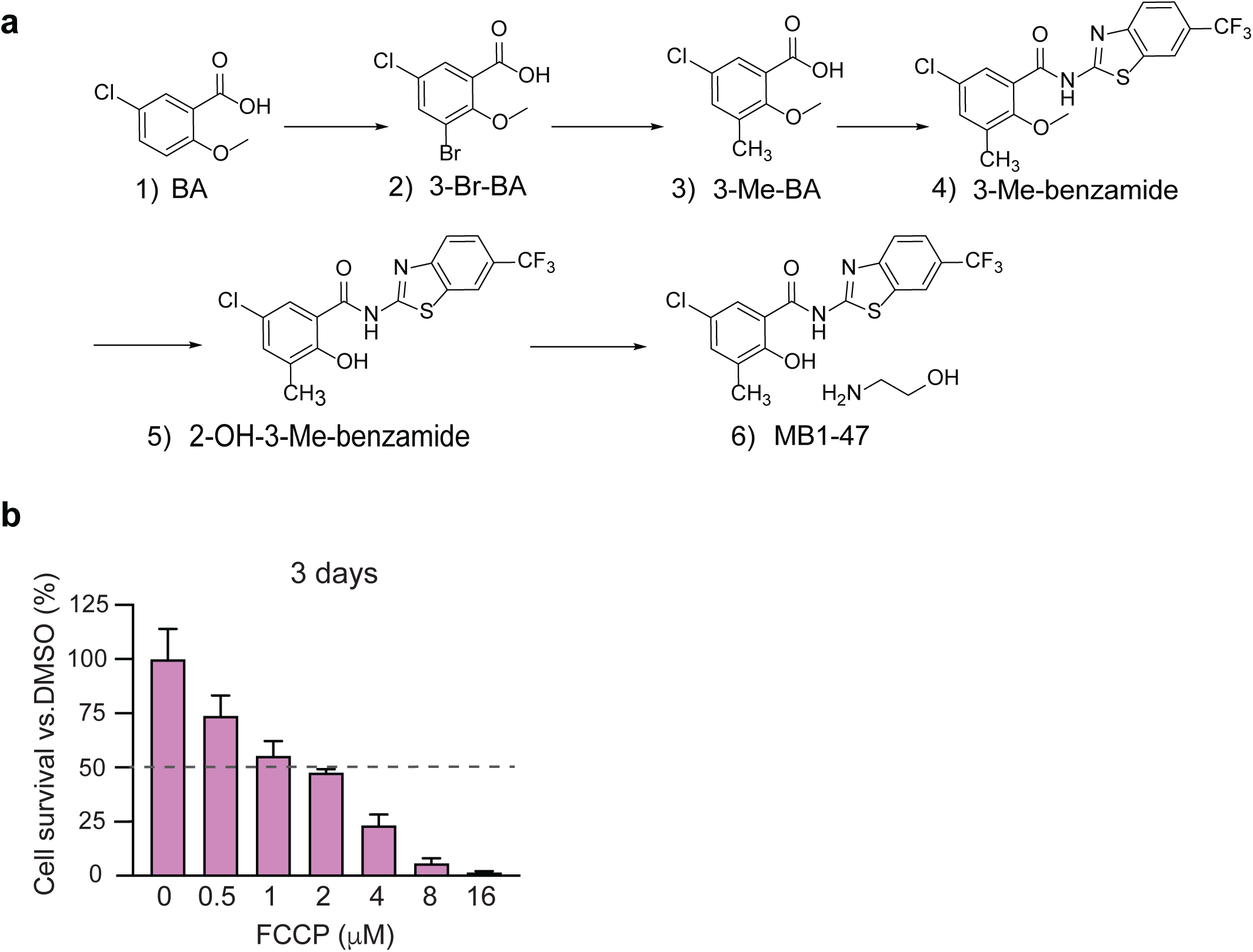
Synthetic route and pharmacological properties of MB1-47. **a**, Synthetic route and chemical structure for MB1-47 (details are described in Methods). **b**, Pharmacokinetic and tissue distribution properties of oral MB1-47 (10 mg/kg). PPB, plasma protein binding; Vss, volume of distribution; Liver/Plasma, distribution ratio between liver (mg/g) and plasma (μg/ml). **c**, Relative cell survival of DND41 cells after 72 h of exposure to the indicated concentrations of FCCP (mean ± s.d.; n=3).

**Extended Fig. 2:**
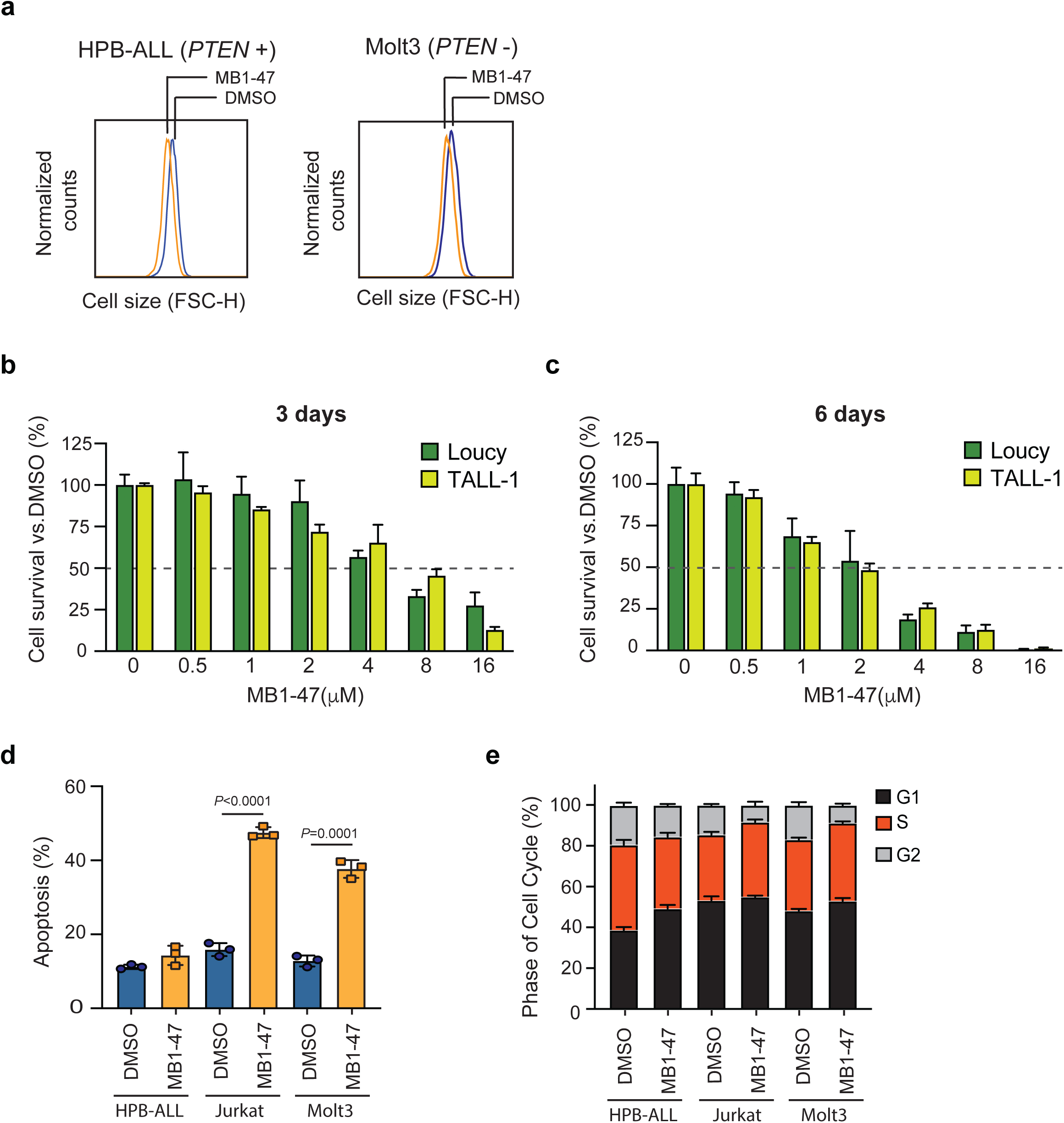
Cytotoxic and cytostatic effects of MB1-47. **a**, Representative flow cytometry histograms showing cell size changes in G1-gated HPB-ALL and Molt3 cells treated in presence or absence of MB1-47 for 72 h. **b-c**, Relative cell survival of Loucy and TALL-1 cells in triplicates in presence of MB1-47 at the indicated concentrations for 3 days (**b)** or 6 days (**c**). **d**, Quantification of apoptotic cells from HPB-ALL, Jurkat and Molt3 cells treated for 72 h with DMSO (control) or MB1-47 (mean ± s.d.; n=3). **e**, Quantification of cell cycle progression in HPB-ALL, Jurkat and Molt3 cells in presence or absence of MB1-47 for 72 h (mean ± s.d.; n=3). Statistical significance (*P*) was determined by using two-tailed Student’s t-test.

**Extended Fig. 3:**
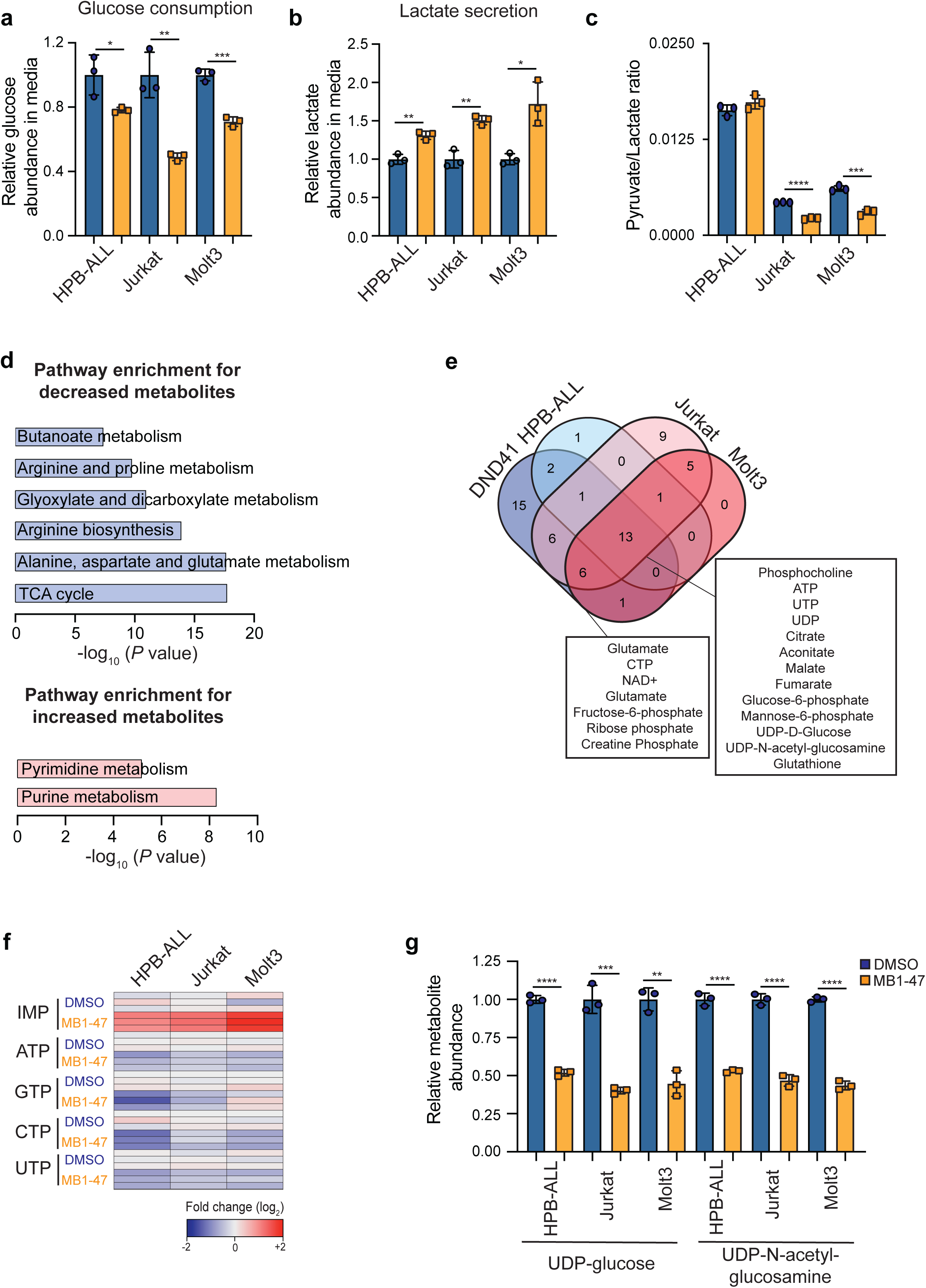
MB1-47 increases glycolysis and compromises nucleotide biosynthesis in T-ALL cell lines. **a**, Relative glucose abundance in media from T-ALL cells cultured in presence or absence of MB1-47 (mean ± s.d.; n = 3). **b**, Relative lactate abundance in media from T-ALL cells cultured in presence or absence of MB1-47 (mean ± s.d.; n = 3). **c**, pyruvate/lactate ratios from indicated T-ALL cells cultured in presence or absence of MB1-47 (mean ± s.d.; n=3). **d**, Pathway enrichment analyses from significantly decreased (top) and increased (bottom) metabolites in DND41 cells after MB1-47 treatment. **e**, Venn diagram showing the common metabolites that significantly drop after MB1-47 exposure in PTEN-positive and PTEN-negative T-ALL cells. **f**, Heat map showing differential intracellular nucleotide abundance (log_2_) after MB1-47 treatment in the indicated T-ALL cells, relative to DMSO-treated controls. **g**, Relative abundance of the indicated UDP-sugars in T-ALL cell lines cultured in presence or absence of MB1-47. All measurements were determined after 24 h of treatment and are relative to DMSO-treated control cells. Statistical significance (*P*) was determined by using two-tailed Student’s t-test. * *P* < 0.05, ** *P* < 0.01, *** *P* < 0.001, **** *P* < 0.0001

**Extended Fig. 4:**
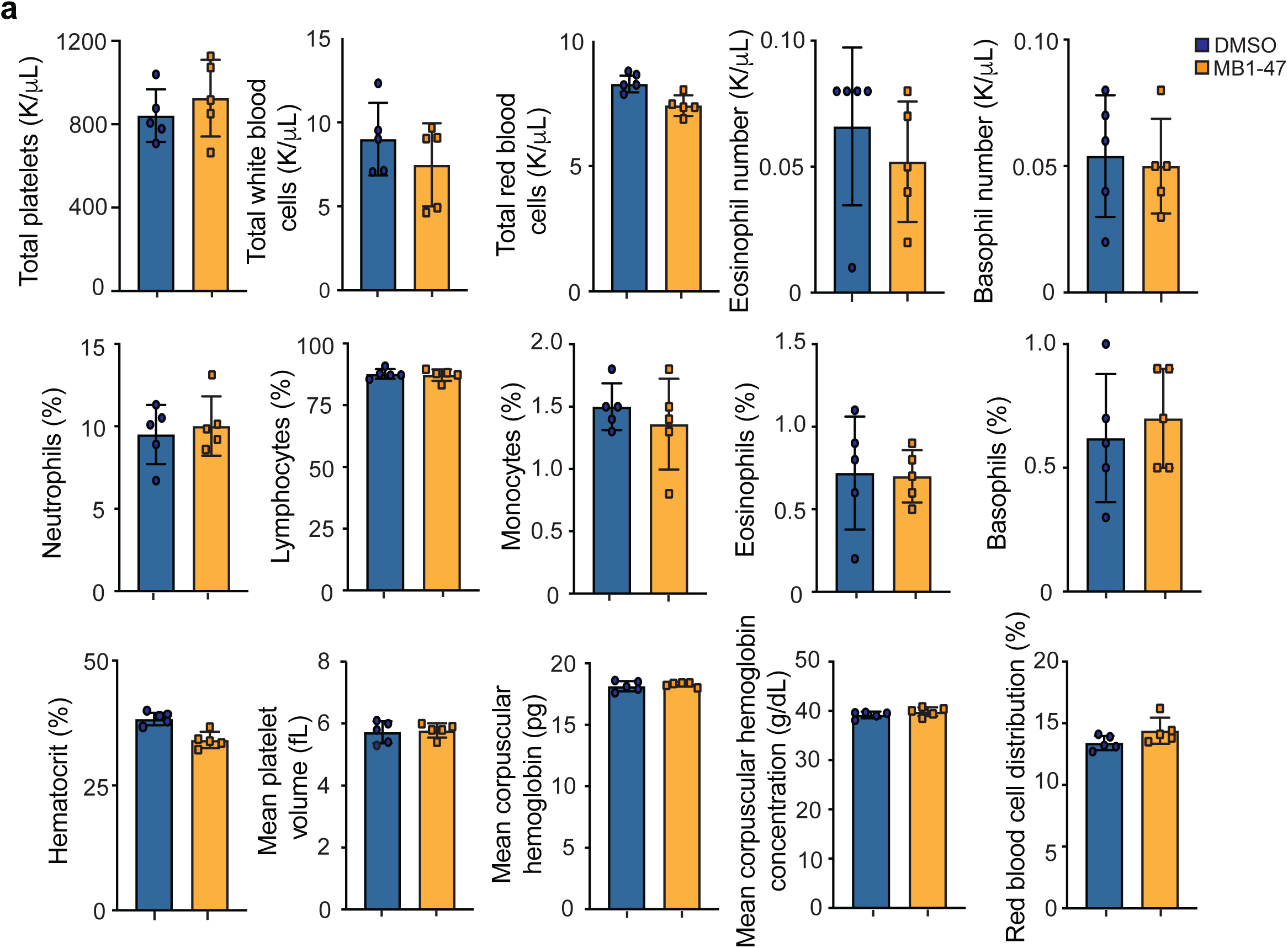
Safety profile of MB1-47. **a**, Hematological parameters of mice fed with control diet or MB1-47-containing diet for 30 consecutive days. Bar graphs represent mean values and error bars represent s.d. Statistical significance (*P*) was determined by using two-tailed Student’s t-test.

## References

1 Litzow, M. R. & Ferrando, A. A. How I treat T-cell acute lymphoblastic leukemia in adults. Blood 126, 833–841, doi:10.1182/blood-2014-10-551895 (2015).

2 Hefazi, M. & Litzow, M. R. Recent advances in the biology and treatment of B-cell acute lymphoblastic leukemia. Blood Lymphat Cancer 8, 47–61, doi:10.2147/BLCTT.S170351 (2018).

3 Hunger, S. P. & Mullighan, C. G. Acute Lymphoblastic Leukemia in Children. N Engl J Med 373, 1541–1552, doi:10.1056/NEJMra1400972 (2015).

4 Kozlowski, P. et al. High relapse rate of T cell acute lymphoblastic leukemia in adults treated with Hyper-CVAD chemotherapy in Sweden. Eur J Haematol 92, 377–381, doi:10.1111/ejh.12269 (2014).

5 Fielding, A. K. et al. Outcome of 609 adults after relapse of acute lymphoblastic leukemia (ALL); an MRC UKALL12/ECOG 2993 study. Blood 109, 944–950, doi:10.1182/blood-2006-05-018192 (2007).

6 Offidani, M., Corvatta, L., Malerba, L., Marconi, M. & Leoni, P. Infectious complications in adult acute lymphoblastic leukemia (ALL): experience at one single center. Leuk Lymphoma 45, 1617–1621, doi:10.1080/10428190410001683660 (2004).

7 Krull, K. R. et al. Neurocognitive outcomes decades after treatment for childhood acute lymphoblastic leukemia: a report from the St Jude lifetime cohort study. J Clin Oncol 31, 4407–4415, doi:10.1200/JCO.2012.48.2315 (2013).

8 Oeffinger, K. C. et al. Chronic health conditions in adult survivors of childhood cancer. N Engl J Med 355, 1572–1582, doi:10.1056/NEJMsa060185 (2006).

9 Hanahan, D. & Weinberg, R. A. Hallmarks of cancer: the next generation. Cell 144, 646–674, doi:10.1016/j.cell.2011.02.013 (2011).

10 Farber, S. & Diamond, L. K. Temporary remissions in acute leukemia in children produced by folic acid antagonist, 4-aminopteroyl-glutamic acid. N Engl J Med 238, 787–793, doi:10.1056/NEJM194806032382301 (1948).

11 Rashkovan, M. & Ferrando, A. Metabolic dependencies and vulnerabilities in leukemia. Genes Dev 33, 1460–1474, doi:10.1101/gad.326470.119 (2019).

12 Garcia-Canaveras, J. C. et al. SHMT inhibition is effective and synergizes with methotrexate in T-cell acute lymphoblastic leukemia. Leukemia, doi:10.1038/s41375-020-0845-6 (2020).

13 Tzoneva, G. et al. Activating mutations in the NT5C2 nucleotidase gene drive chemotherapy resistance in relapsed ALL. Nat Med 19, 368–371, doi:10.1038/nm.3078 (2013).

14 Tzoneva, G. et al. Clonal evolution mechanisms in NT5C2 mutant-relapsed acute lymphoblastic leukaemia. Nature 553, 511–514, doi:10.1038/nature25186 (2018).

15 Calvert, H. Folate status and the safety profile of antifolates. Semin Oncol 29, 3–7, doi:10.1016/s0093-7754(02)70209-1 (2002).

16 Genestier, L. et al. Immunosuppressive properties of methotrexate: apoptosis and clonal deletion of activated peripheral T cells. J Clin Invest 102, 322–328, doi:10.1172/JCI2676 (1998).

17 Ohnuma, T., Holland, J. F., Freeman, A. & Sinks, L. F. Biochemical and pharmacological studies with asparaginase in man. Cancer Res 30, 2297–2305 (1970).

18 Boyse, E. A., Old, L. J., Campbell, H. A. & Mashburn, L. T. Suppression of murine leukemias by L-asparaginase. Incidence of sensitivity among leukemias of various types: comparative inhibitory activities of guinea pig serum L-asparaginase and Escherichia coli L-asparaginase. J Exp Med 125, 17–31, doi:10.1084/jem.125.1.17 (1967).

19 Horowitz, B. et al. Asparagine synthetase activity of mouse leukemias. Science 160, 533–535, doi:10.1126/science.160.3827.533 (1968).

20 Hijiya, N. & van der Sluis, I. M. Asparaginase-associated toxicity in children with acute lymphoblastic leukemia. Leuk Lymphoma 57, 748–757, doi:10.3109/10428194.2015.1101098 (2016).

21 Weng, A. P. et al. Activating mutations of NOTCH1 in human T cell acute lymphoblastic leukemia. Science 306, 269–271, doi:10.1126/science.1102160 (2004).

22 Selkoe, D. & Kopan, R. Notch and Presenilin: regulated intramembrane proteolysis links development and degeneration. Annu Rev Neurosci 26, 565–597, doi:10.1146/annurev.neuro.26.041002.131334 (2003).

23 Wei, P. et al. Evaluation of selective gamma-secretase inhibitor PF-03084014 for its antitumor efficacy and gastrointestinal safety to guide optimal clinical trial design. Mol Cancer Ther 9, 1618–1628, doi:10.1158/1535-7163.MCT-10-0034 (2010).

24 Palomero, T. et al. Mutational loss of PTEN induces resistance to NOTCH1 inhibition in T-cell leukemia. Nat Med 13, 1203–1210, doi:10.1038/nm1636 (2007).

25 Kishton, R. J. et al. AMPK Is Essential to Balance Glycolysis and Mitochondrial Metabolism to Control T-ALL Cell Stress and Survival. Cell Metab 23, 649–662, doi:10.1016/j.cmet.2016.03.008 (2016).

26 Herranz, D. et al. Metabolic reprogramming induces resistance to anti-NOTCH1 therapies in T cell acute lymphoblastic leukemia. Nat Med 21, 1182–1189, doi:10.1038/nm.3955 (2015).

27 Palomero, T. et al. CUTLL1, a novel human T-cell lymphoma cell line with t(7;9) rearrangement, aberrant NOTCH1 activation and high sensitivity to gamma-secretase inhibitors. Leukemia 20, 1279–1287, doi:10.1038/sj.leu.2404258 (2006).

28 Melamud, E., Vastag, L. & Rabinowitz, J. D. Metabolomic analysis and visualization engine for LC-MS data. Anal Chem 82, 9818–9826, doi:10.1021/ac1021166 (2010).

29 Su, X., Lu, W. & Rabinowitz, J. D. Metabolite Spectral Accuracy on Orbitraps. Anal Chem 89, 5940–5948, doi:10.1021/acs.analchem.7b00396 (2017).

30 Antoniewicz, M. R., Kelleher, J. K. & Stephanopoulos, G. Elementary metabolite units (EMU): a novel framework for modeling isotopic distributions. Metab Eng 9, 68–86, doi:10.1016/j.ymben.2006.09.001 (2007).

31 DeBerardinis, R. J. & Chandel, N. S. Fundamentals of cancer metabolism. Sci Adv 2, e1600200, doi:10.1126/sciadv.1600200 (2016).

32 Chen, W., Mook, R. A., Jr., Premont, R. T. & Wang, J. Niclosamide: Beyond an antihelminthic drug. Cell Signal 41, 89–96, doi:10.1016/j.cellsig.2017.04.001 (2018).

33 Li, Y. et al. Multi-targeted therapy of cancer by niclosamide: A new application for an old drug. Cancer Lett 349, 8–14, doi:10.1016/j.canlet.2014.04.003 (2014).

34 Schweizer, M. T. et al. A phase I study of niclosamide in combination with enzalutamide in men with castration-resistant prostate cancer. PLoS One 13, e0198389, doi:10.1371/journal.pone.0198389 (2018).

35 Burock, S. et al. Phase II trial to investigate the safety and efficacy of orally applied niclosamide in patients with metachronous or sychronous metastases of a colorectal cancer progressing after therapy: the NIKOLO trial. BMC Cancer 18, 297, doi:10.1186/s12885-018-4197-9 (2018).

36 Maragos, W. F. & Korde, A. S. Mitochondrial uncoupling as a potential therapeutic target in acute central nervous system injury. J Neurochem 91, 257–262, doi:10.1111/j.1471-4159.2004.02736.x (2004).

37 Amara, C. E. et al. Mild mitochondrial uncoupling impacts cellular aging in human muscles in vivo. Proc Natl Acad Sci U S A 104, 1057–1062, doi:10.1073/pnas.0610131104 (2007).

38 Chen, B. et al. Computational Discovery of Niclosamide Ethanolamine, a Repurposed Drug Candidate That Reduces Growth of Hepatocellular Carcinoma Cells In Vitro and in Mice by Inhibiting Cell Division Cycle 37 Signaling. Gastroenterology 152, 2022–2036, doi:10.1053/j.gastro.2017.02.039 (2017).

39 Alasadi, A. et al. Effect of mitochondrial uncouplers niclosamide ethanolamine (NEN) and oxyclozanide on hepatic metastasis of colon cancer. Cell Death Dis 9, 215, doi:10.1038/s41419-017-0092-6 (2018).

40 Piovan, E. et al. Direct reversal of glucocorticoid resistance by AKT inhibition in acute lymphoblastic leukemia. Cancer Cell 24, 766–776, doi:10.1016/j.ccr.2013.10.022 (2013).

41 Samudio, I., Fiegl, M., McQueen, T., Clise-Dwyer, K. & Andreeff, M. The warburg effect in leukemia-stroma cocultures is mediated by mitochondrial uncoupling associated with uncoupling protein 2 activation. Cancer Res 68, 5198–5205, doi:10.1158/0008-5472.CAN-08-0555 (2008).

42 Griss, T. et al. Metformin Antagonizes Cancer Cell Proliferation by Suppressing Mitochondrial-Dependent Biosynthesis. PLoS Biol 13, e1002309, doi:10.1371/journal.pbio.1002309 (2015).

43 Sullivan, L. B. et al. Supporting Aspartate Biosynthesis Is an Essential Function of Respiration in Proliferating Cells. Cell 162, 552–563, doi:10.1016/j.cell.2015.07.017 (2015).

44 Birsoy, K. et al. An Essential Role of the Mitochondrial Electron Transport Chain in Cell Proliferation Is to Enable Aspartate Synthesis. Cell 162, 540–551, doi:10.1016/j.cell.2015.07.016 (2015).

45 Chouchani, E. T. et al. Ischaemic accumulation of succinate controls reperfusion injury through mitochondrial ROS. Nature 515, 431–435, doi:10.1038/nature13909 (2014).

46 Pollyea, D. A. et al. Venetoclax with azacitidine disrupts energy metabolism and targets leukemia stem cells in patients with acute myeloid leukemia. Nat Med 24, 1859–1866, doi:10.1038/s41591-018-0233-1 (2018).

47 Sullivan, L. B. et al. Aspartate is an endogenous metabolic limitation for tumour growth. Nat Cell Biol 20, 782–788, doi:10.1038/s41556-018-0125-0 (2018).

48 Garcia-Bermudez, J. et al. Aspartate is a limiting metabolite for cancer cell proliferation under hypoxia and in tumours. Nat Cell Biol 20, 775–781, doi:10.1038/s41556-018-0118-z (2018).

49 Mullen, A. R. et al. Reductive carboxylation supports growth in tumour cells with defective mitochondria. Nature 481, 385–388, doi:10.1038/nature10642 (2011).

50 Shackelford, D. B. & Shaw, R. J. The LKB1-AMPK pathway: metabolism and growth control in tumour suppression. Nat Rev Cancer 9, 563–575, doi:10.1038/nrc2676 (2009).

51 Chan, S. M., Weng, A. P., Tibshirani, R., Aster, J. C. & Utz, P. J. Notch signals positively regulate activity of the mTOR pathway in T-cell acute lymphoblastic leukemia. Blood 110, 278–286, doi:10.1182/blood-2006-08-039883 (2007).

52 Lee, K. et al. Vital roles of mTOR complex 2 in Notch-driven thymocyte differentiation and leukemia. J Exp Med 209, 713–728, doi:10.1084/jem.20111470 (2012).

53 Belver, L. & Ferrando, A. The genetics and mechanisms of T cell acute lymphoblastic leukaemia. Nat Rev Cancer 16, 494–507, doi:10.1038/nrc.2016.63 (2016).

54 Weinberg, F. et al. Mitochondrial metabolism and ROS generation are essential for Kras-mediated tumorigenicity. Proc Natl Acad Sci U S A 107, 8788–8793, doi:10.1073/pnas.1003428107 (2010).

55 Guo, J. Y. et al. Activated Ras requires autophagy to maintain oxidative metabolism and tumorigenesis. Genes Dev 25, 460–470, doi:10.1101/gad.2016311 (2011).

56 Marin-Valencia, I. et al. Analysis of tumor metabolism reveals mitochondrial glucose oxidation in genetically diverse human glioblastomas in the mouse brain in vivo. Cell Metab 15, 827–837, doi:10.1016/j.cmet.2012.05.001 (2012).

57 Wheaton, W. W. et al. Metformin inhibits mitochondrial complex I of cancer cells to reduce tumorigenesis. Elife 3, e02242, doi:10.7554/eLife.02242 (2014).

58 Molina, J. R. et al. An inhibitor of oxidative phosphorylation exploits cancer vulnerability. Nat Med 24, 1036–1046, doi:10.1038/s41591-018-0052-4 (2018).

59 Ibrahim, A., Yucel, N., Kim, B. & Arany, Z. Local Mitochondrial ATP Production Regulates Endothelial Fatty Acid Uptake and Transport. Cell Metab, doi:10.1016/j.cmet.2020.05.018 (2020).

60 Carrer, A. & Wellen, K. E. Metabolism and epigenetics: a link cancer cells exploit. Curr Opin Biotechnol 34, 23–29, doi:10.1016/j.copbio.2014.11.012 (2015).

61 Pietrocola, F., Galluzzi, L., Bravo-San Pedro, J. M., Madeo, F. & Kroemer, G. Acetyl coenzyme A: a central metabolite and second messenger. Cell Metab 21, 805–821, doi:10.1016/j.cmet.2015.05.014 (2015).

62 Vara-Ciruelos, D. et al. Phenformin, But Not Metformin, Delays Development of T Cell Acute Lymphoblastic Leukemia/Lymphoma via Cell-Autonomous AMPK Activation. Cell Rep 27, 690–698 e694, doi:10.1016/j.celrep.2019.03.067 (2019).

